# Downregulation of NR4A3 during inactivity alters glucose metabolism and impairs translation in human skeletal muscle

**DOI:** 10.1101/2024.03.11.584422

**Authors:** Jonathon A.B. Smith, Brendan M. Gabriel, Ahmed M. Abdelmoez, Mladen Savikj, Shane C. Wright, Stefania Koutsilieri, Romain Barrès, Volker M. Lauschke, Anna Krook, Juleen R. Zierath, Nicolas J. Pillon

## Abstract

The transcription factor NR4A3 is regulated by exercise and a potent modulator of skeletal muscle metabolism. We report here that physical inactivity decreased skeletal muscle NR4A3 levels, concomitant with mitochondrial function and proteostasis pathways. Silencing of NR4A3 in myotubes decreased glucose oxidation and increased lactate production. This coincided with greater signalling downstream of AMPK and elevated rates of fatty acid oxidation. While NR4A3 silencing reduced protein synthesis, mTORC1 signalling, and ribosomal transcription, overexpression of the canonical NR4A3 protein isoform augmented translation and cellular protein content. Moreover, restoration of NR4A3 levels rescued glucose oxidation in NR4A3-silenced myotubes and restored phosphorylation of mTORC1 substrates. NR4A3 depletion reduced myotube area and altered the abundance of contractile elements. Thus, downregulation of NR4A3 has adverse effects on skeletal muscle metabolism, myotube size, and contractile apparatus by directing mTORC1 signalling and ribosomal biogenesis. Our data demonstrate that NR4A3 controls skeletal muscle atrophy associated with physical inactivity.

## Introduction

Engagement in physical activity initiates a cascade of integrative responses related to skeletal muscle contraction, such as increased energy expenditure, haemodynamics, and oxygen consumption (Smith *et al*, 2023). Depending on the modality, intensity, and volume of exercise performed, these homeostatic processes trigger a myriad of beneficial downstream events that serve to maintain or enhance aspects of human physiology, including cardiovascular function, skeletal muscle mass, and peripheral insulin sensitivity (Smith *et al*., 2023). Conversely, inactivity impairs skeletal muscle size and composition. These alterations have adverse effects on local tissue function and metabolism, which in turn impacts systemic health. Inactivity diminishes physical performance (Brook *et al*, 2022; Manini *et al*, 2007), whole-body metabolic flexibility (Bergouignan *et al*, 2013), and is associated with reduced quality of life (Yerrakalva *et al*, 2023) and increased risk of non-communicable diseases like type 2 diabetes (Yuan *et al*, 2023).

Changes in skeletal muscle mass are determined by rates of myofibrillar protein synthesis versus degradation. Physical inactivity favours signalling and transcriptional programmes that promote sustained periods of negative protein balance and muscle loss. Indeed, detectable muscle atrophy occurs within just four days of disuse (Brook *et al*., 2022; Shur *et al*, 2024). One of the major regulators of translation is the mammalian target of rapamycin complex 1 (mTORC1), which permits myofibrillar protein synthesis and muscle hypertrophy (You *et al*, 2019). However, mTORC1 acts in tandem with ribosomal content (among other processes) to regulate translation (West *et al*, 2016). Detriments in both mTORC1 (Shad *et al*, 2019) and ribosomal abundance (Figueiredo *et al*, 2021) are observed in skeletal muscle with inactivity. Alongside insulin- and amino acid-resistance (Breen *et al*, 2013; Brook *et al*., 2022; Pavis *et al*, 2023; Shad *et al*., 2019; Shur *et al*., 2024), this downregulation of the translational machinery drives initial losses in skeletal muscle mass through dampened myofibrillar protein synthetic responses (Brook *et al*., 2022; Pavis *et al*., 2023). As an effector of the phosphatidylinositol-3-kinase (PI3K)-AKT pathway, mTORC1 is also central to proteostasis via temporal coordination of anabolic and catabolic processes (Kaiser *et al*, 2022). Nevertheless, the molecular transducers of this inactivity phenotype have not been fully elucidated.

Exercise training and physical inactivity are not mere opposing ends of a linear spectrum, but rather multifaceted processes with distinct underlying mechanisms. Understanding this dichotomy between exercise and inactivity necessitates recognising that they engage both common and distinct signalling and transcriptional pathways (Pillon *et al*, 2020). The NR4A family of stress-responsive orphan nuclear receptors are commonly induced by exercise and all members have ties to skeletal muscle metabolism, development, or remodelling. However, only nuclear receptor subfamily 4 group A member 3 (*NR4A3*, also known as *NOR1*) shows opposite regulation to exercise and inactivity (Pillon *et al*., 2020). Depletion of NR4A3 in primary human (Pillon *et al*., 2020) or mouse C2C12 (Paez *et al*, 2023) skeletal muscle myotubes reduces mitochondrial oxidative capacity. Alternatively, transgenic overexpression of *Nr4a3* (Pearen *et al*, 2012) or *Nr4a1* (also known as *Nur77*) (Chao *et al*, 2012) shifts skeletal muscle composition towards an oxidative phenotype, suggesting functional redundancy between these NR4A family members through shared NGFI-B (Wilson *et al*, 1991) or Nur (Maira *et al*, 1999) response elements. Furthermore, overexpression of *Nr4a3* (Pearen *et al*., 2012), *Nr4a1* (Chao *et al*., 2012), or *Nr4a2* (also known as *Nurr1*) (Amoasii *et al*, 2019) in skeletal muscle enhances physical performance, while deletion of *Nr4a1* impairs muscle size (Tontonoz *et al*, 2015). Silencing of *Nr4a3* in mouse C2C12 myotubes reduces mTORC1 signalling and global myosin heavy chain gene expression (Paez *et al*., 2023) but the molecular mechanisms underlying the metabolic, anabolic, and transcriptomic roles of NR4A3 remain largely unknown in human skeletal muscle.

Here, we investigated the impact of NR4A3 on metabolism and post-mitotic growth processes. We provide evidence that *NR4A3* is reduced during skeletal muscle disuse and that downregulation of *NR4A3* impairs glucose metabolism and protein synthesis, contributing towards atrophy of skeletal muscle cells.

## Results and Discussion

### Downregulation of *NR4A3* during inactivity is associated with remodelling of myogenic and metabolic pathways

*NR4A3* mRNA is induced by exercise and repressed during inactivity in human skeletal muscle (Pillon *et al*., 2020). We previously reported that gene expression profiles of primary human skeletal myotubes subjected to *NR4A3* silencing best reflects the human muscle transcriptome after periods of bedrest and limb immobilisation (Pillon *et al*., 2020). To build upon this observation and gain further insight into the regulation of *NR4A3*, we analysed eight published transcriptomic studies of the human skeletal muscle response to inactivity. The meta-analysis revealed that physical inactivity reduced *NR4A3* mRNA by 27% (95% CI: 42, 9), but this decrease was not evident in all studies (**Figure 1A**). Grouping studies based on the duration of inactivity demonstrated that skeletal muscle *NR4A3* expression is transiently repressed during the first days of disuse before returning to baseline levels after 2 weeks (**Figure 1B**). Correlation of *NR4A3* with all other transcripts present in the eight studies allowed for the identification of pathways co-regulated with *NR4A3* during inactivity (**Figure 1C**). Gene set enrichment analysis showed that *NR4A3* was positively associated with mitochondrial function and negatively associated with pathways related to cytoskeleton organisation, chromatin regulation, protein synthesis, and degradation (**Figure 1C**). Furthermore, combined analysis of two datasets exploring the restoration of ambulatory behaviour after inactivity showed that reloading re-established *NR4A3* mRNA to pre-inactivity levels (**Figure 1D**), consistent with its role as a contraction-responsive transcription factor in skeletal muscle (Pattamaprapanont *et al*, 2016; Pillon *et al*., 2020).

**Figure 1.**
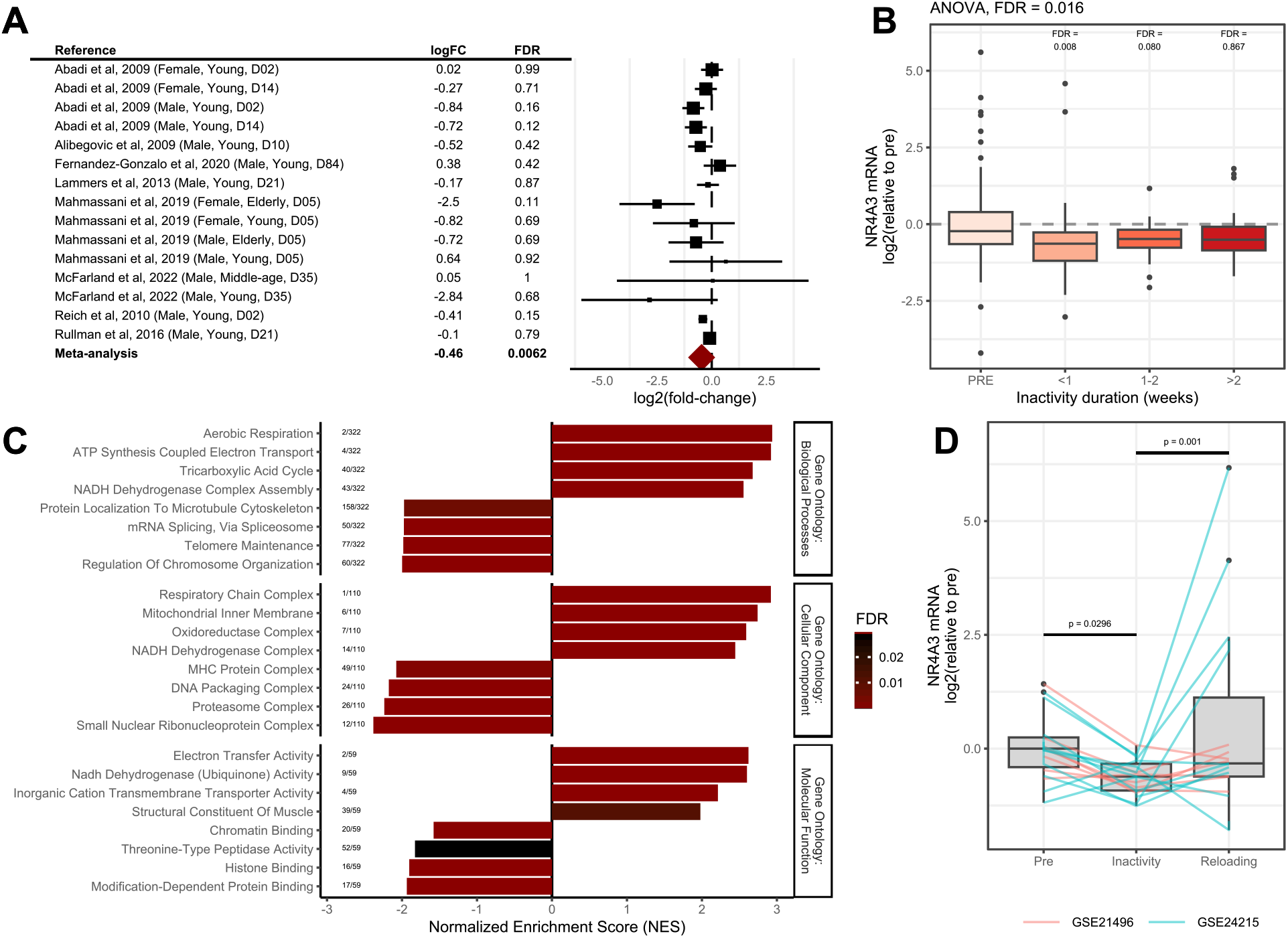
Downregulation of *NR4A3* during inactivity is associated with remodelling of myogenic and metabolic pathways. **A**. Meta-analysis of transcriptomic inactivity studies in human skeletal muscle.**B**. Transcriptomic studies were pooled by the duration of inactivity protocols: less than one week, one-two weeks or more than two weeks. Datasets were normalized as described in methods. **C**. Gene set enrichment analysis with the Gene Ontology dataset was performed on genes ranked on Spearman correlation with *NR4A3* across all transcriptomic studies of inactivity. Bar colour represents FDR and numbers to the left of each bar indicate the rank of each ontology within the total number of enriched pathways. **D**. The two studies that explored effects of reloading after inactivity were merged and analysed as described in methods.

### Silencing *NR4A3* attenuates glucose oxidation by diverting glucose towards lactate and increases fatty acid oxidation

To mimic the decreased levels observed in human skeletal muscle after inactivity, *NR4A3* was experimentally downregulated using RNA interference in differentiated primary human skeletal muscle cells. Silencing of *NR4A3* was associated with modest compensatory increases in other NR4A family members *NR4A1* (*NUR77*) and *NR4A2* (*NURR1*) (**Figure 2A**). NR4A3 protein was predominantly localised to the nucleus in myotubes and silencing efficiently reduced nuclear protein abundance (**Figure 2B**). Importantly, this depletion of NR4A3 lowered basal and FCCP-stimulated (uncoupled) glucose oxidation (**Figure 2C**) independent of glucose transport. However, several phosphorylation events in the canonical insulin signalling cascade were potentiated by silencing of *NR4A3* (**Supplementary Figure 1**), including site Thr308 on AKT and downstream inactivating phosphorylation of both AKT substrate of 160 kDa (AS160, also known as TBC1D4) and glycogen synthase kinase 3 (GSK3), which facilitate glucose transporter 4 (GLUT4) translocation and glycogen storage, respectively. Furthermore, basal and insulin-mediated glucose uptake and incorporation into glycogen responses were maintained (**Supplementary Figures 1H, 1I**).

**Figure 2.**
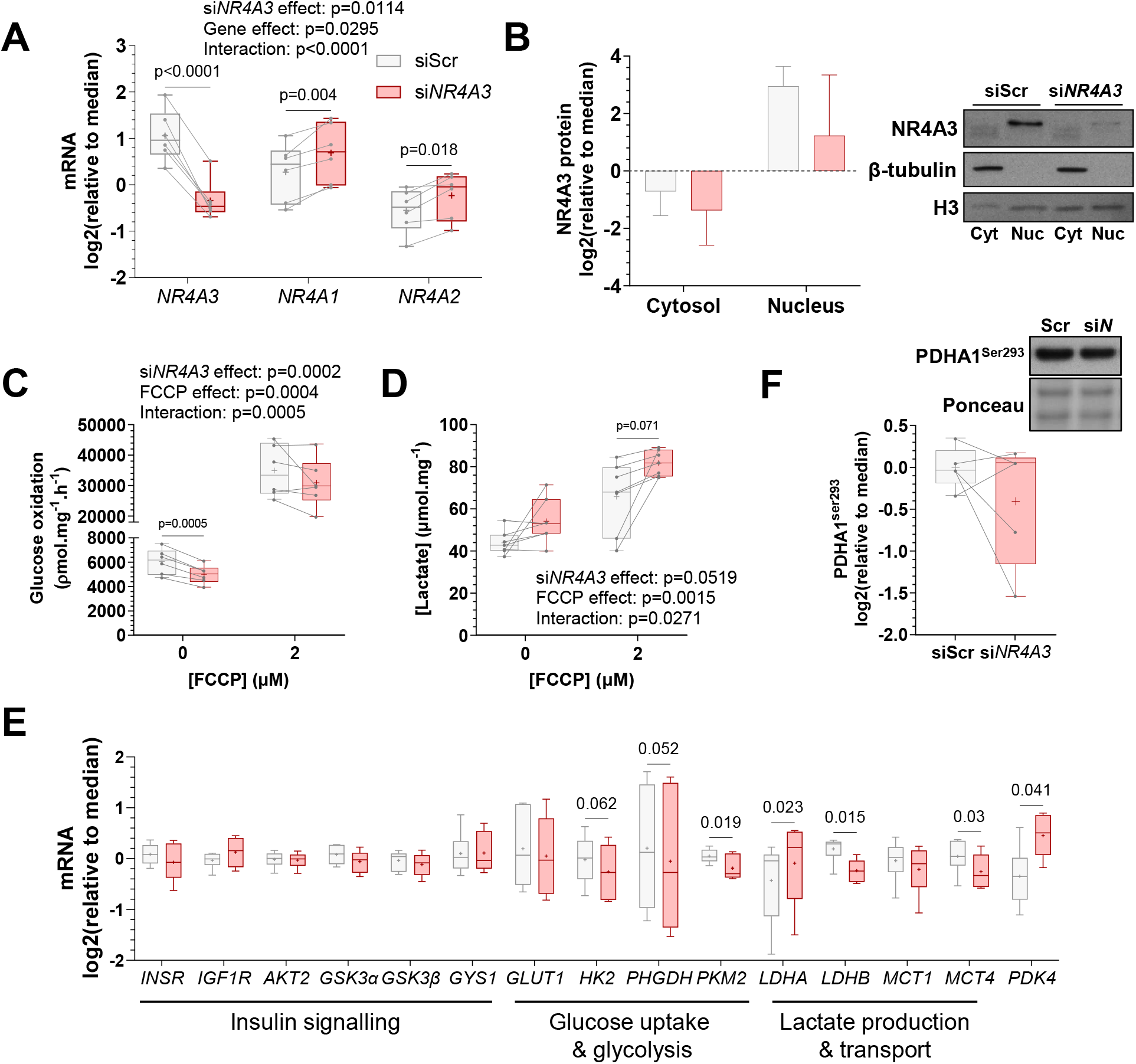
*NR4A3* silencing in primary human myotubes attenuates glucose oxidation by diverting glucose towards lactate production. Primary skeletal muscle cells were exposed to a control scramble sequence (siScr) or a silencing RNA targeting *NR4A3* (si*NR4A3*). **A**. mRNA expression of NR4A family members *NR4A3, NR4A1* and *NR4A2* measured by RT-qPCR. n = 6, 2-way ANOVA (silencing x gene) with Šidák correction. **B**. Protein level of NR4A3 in cytosolic and nuclear fractions assessed by immunoblot analysis. Results are mean ± SEM, n = 2. **C**. Rates of radiolabelled glucose oxidation under basal and (2 µM) FCCP-stimulated conditions over 4 h. n = 6, 2-way ANOVA (silencing x FCCP) with Šidák correction. **D**. Lactate concentration in cell supernatant measured after 48 h of basal or (2 µM) FCCP-stimulated conditions. n = 7, 2-way ANOVA (silencing x FCCP) with Šidák correction. Overall statistical model effects are stated in figures. **E**. mRNA expression of genes involved in glucose metabolism measured by RT-qPCR. Results are box-and-whisker plots with Tukey distribution and crosses indicating mean values. n = 6, paired t-tests with FDR correction. **F**. Representative immunoblot and quantification of pyruvate dehydrogenase E1 Subunit Alpha 1 (PDHA1) phosphorylation at Ser^293^. n = 5, 2-way ANOVA (silencing x insulin + leucine).

Upon *NR4A3* silencing, lactate production and release into culture medium was augmented (**Figure 2D**), indicating a shift in glucose fate that was also reported in mouse C2C12 myotubes (Pearen *et al*, 2008). Accordingly, mRNA expression of the alpha isoform of lactate dehydrogenase (*LDHA*) increased after *NR4A3* silencing, while expression of the beta isoform decreased (*LDHB*) (**Figure 2E**). While total LDH protein abundance remained unaltered (**Supplementary Figure 1K**), the modification of LDH isoform composition indicates preferential production of lactate from glycolysis and/or less conversion of lactate to pyruvate (Liang *et al*, 2016). This notion was further supported by the reduction of pyruvate kinase muscle 2 (*PKM2*) mRNA (**Figure 2E**), as inhibiting noncanonical PKM2 activity increased lactate release from C2C12 myoblasts (Blum *et al*, 2021). For glucose to be oxidised in the mitochondria, pyruvate must be metabolised to acetyl coenzyme A (acetyl-CoA) via the pyruvate dehydrogenase (PDH) complex. *NR4A3* silencing upregulated pyruvate dehydrogenase kinase 4 (*PDK4*) gene expression (**Figure 2E**) consistent with attenuated glucose oxidation (Pin *et al*, 2019) (**Figure 2C**). However, the inhibitory PDK4-target phosphorylation site on PDH E1 component subunit alpha (PDHA1^Ser293^) was unaltered (**Figure 2F**), indicating that PDH activity was not perturbed.

We previously reported that mitochondrial respiration (in the presence of glucose, glutamine, and pyruvate) and protein subunits of the electron transport chain were diminished by *NR4A3* silencing in primary human skeletal myotubes (Pillon *et al*., 2020). This aligns with findings in C2C12 myotubes, where *Nr4a3* depletion decreased mitochondrial membrane potential in the absence of changes in mitochondrial DNA (Paez *et al*., 2023). Instead, downregulation of *Nr4a3* negatively affected the expression of respiratory complex genes, as well as transcripts for mitochondrial ribosomal proteins (Paez *et al*., 2023). Additionally, *Nr4a3* silencing reduced the mRNA abundance of mitofusin-2 (*Mfn2*) but increased dynamin 1 like protein expression (DNM1L; also known as DRP1) (Paez *et al*., 2023). These data suggest that loss of NR4A3 attenuates mitochondrial fusion dynamics (instead favouring fission), mitochondrial reticulum connectivity, and assembly of electron transport chain complexes in a manner separate from alterations in mitochondrial abundance. Our finding that myotube’s response to the mitochondrial uncoupler FCCP was maintained after *NR4A3* RNA interference further supports not only retained mitochondrial content, but also the presence of functional mitochondria (**Figure 2C)**. Thus, the attenuation of glucose oxidation in NR4A3-depleted myotubes appears to be a consequence of greater diversion of glucose towards lactate, which would reduce glucose-derived acetyl-CoA entry into the tricarboxylic acid (TCA) cycle.

Skeletal muscle cells are highly glycolytic in culture, preferentially oxidising glucose at rates 100-fold faster than long-chain lipids (Abdelmoez *et al*, 2020). Therefore, we explored the consequence of NR4A3-dependent reductions in glucose oxidation on lipid metabolism in primary human skeletal myotubes. NR4A3 depletion upregulated basal and FCCP-stimulated rates of fatty acid oxidation (**Figure 3A**) without altering the intracellular composition of select lipid species. Whereas the ratio of triacylglycerides, diacylglycerides, free fatty acids, and other lipids were modified by FCCP, *NR4A3* RNA interference was without effect (**Figure 3B**). Compatible with greater fatty acid oxidation, *NR4A3* silencing increased phosphorylation of the AMP-activated protein kinase α subunit (AMPKα^Thr172^) and its substrates acetyl-CoA carboxylase (ACC^Ser79^) and TBC domain family member 1 (TBC1D1^Ser237^) (**Figure 3C**). This signalling coalesced with activating phosphorylation of hormone-sensitive lipase (HSL^Ser660^), reduced LIPIN1 protein abundance (**Figure 3D**), and downregulation of total ACC protein (**Figure 3C**), as well as the mRNA of its cytosolic isoform (*ACCα*) (**Figure 3E**). Further profiling of transcripts involved in fatty acid metabolic pathways revealed that NR4A3 depletion altered the composition of *AMPKα* isoforms and fatty acid transport genes (**Figure 3E**). Compared to control, *NR4A3* silencing increased the expression of fatty acid transport protein 1 (*FATP1*) and reduced the expression of fatty acid binding protein 3 (*FABP3*) (**Figure 3E**). Together, the protein and transcriptional changes induced after RNA interference of *NR4A3* suggest a physiological mechanism compensating for energy stress due to impaired glucose oxidation.

**Figure 3.**
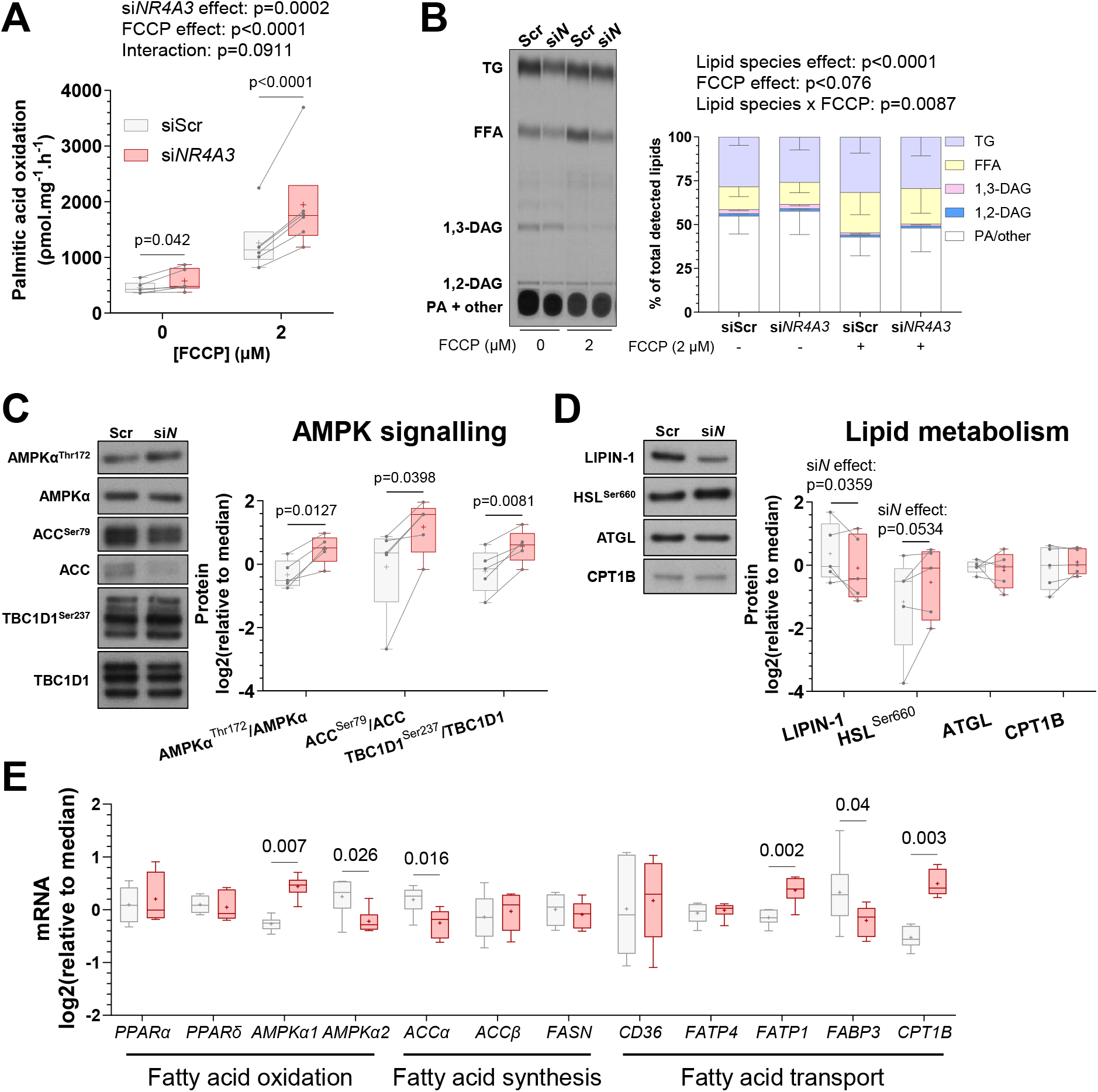
*NR4A3* silencing upregulates fatty acid oxidation and AMPK signalling. Primary skeletal muscle cells were exposed to a control scramble sequence (siScr) or a silencing RNA targeting *NR4A3* (si*NR4A3*). **A**. Rates of radiolabelled ^14^C palmitic acid oxidation under basal and (2 µM) FCCP-stimulated conditions over 4 h. n = 6, 2-way ANOVA (silencing x FCCP) with Šidák correction. **B**. Detection of select lipid species by thin layer chromatography (TLC) of lipid extracts from myotubes after 6 h of basal or (2 µM) FCCP-stimulated conditions. TG = triacylglycerides, FFA = free fatty acids, DAG = diacylglycerol, PA + other = phosphatidic acid plus all other non-migrating lipid species. Results are the mean ± SEM of each lipid species as a percentage of total detected lipid species within condition. n = 5, 3-way ANOVA (silencing x lipid species x FCCP). Overall statistical model effects are stated in figures. **C**. Immunoblot of AMPK-signalling proteins from a representative donor and quantification of phosphorylated-to-total protein ratios. n = 5, 2-way ANOVA (silencing x insulin + leucine) with Šidák correction. Only basal data are shown. **D**. Representative immunoblot and quantification of proteins involved in lipid storage, mobilisation, and transport. n = 5, 2-way ANOVA (silencing x insulin + leucine) with Šidák correction. Only basal data are shown and overall effects of silencing in the model are presented (si*N* effect, p<0.1). **E**. mRNA expression of genes involved in lipid metabolism measured by RT-qPCR. Results are box-and-whisker plots with Tukey distribution and crosses indicating mean values. n = 6, paired t-tests with FDR correction.

### *NR4A3* silencing in primary human myotubes downregulates translation, mTORC1 signalling, and pre-rRNA abundance

Metabolic assays are commonly normalised to protein content to account for differences in seeding density. We observed that silencing of *NR4A3* consistently reduced the total protein and RNA abundance of cultured human myotubes (**Figures 4A, 4B**). These findings, combined with the association of reduced *NR4A3* levels during bed rest and limb immobilisation inactivity (**Figure 1**), led us to explore the putative effects of *NR4A3* silencing on protein synthesis. Relative rates of translation can be measured *in vitro* by the surface sensing of translation (SUnSET) method, which detects puromycin incorporation into nascent peptide chains by immunoblot analysis (Goodman *et al*, 2011). Using this technique, we observed reduced puromycilation of proteins upon *NR4A3* silencing both at baseline and after insulin plus leucine stimulation (**Figure 4C**), suggesting impaired protein synthesis with NR4A3 downregulation.

**Figure 4.**
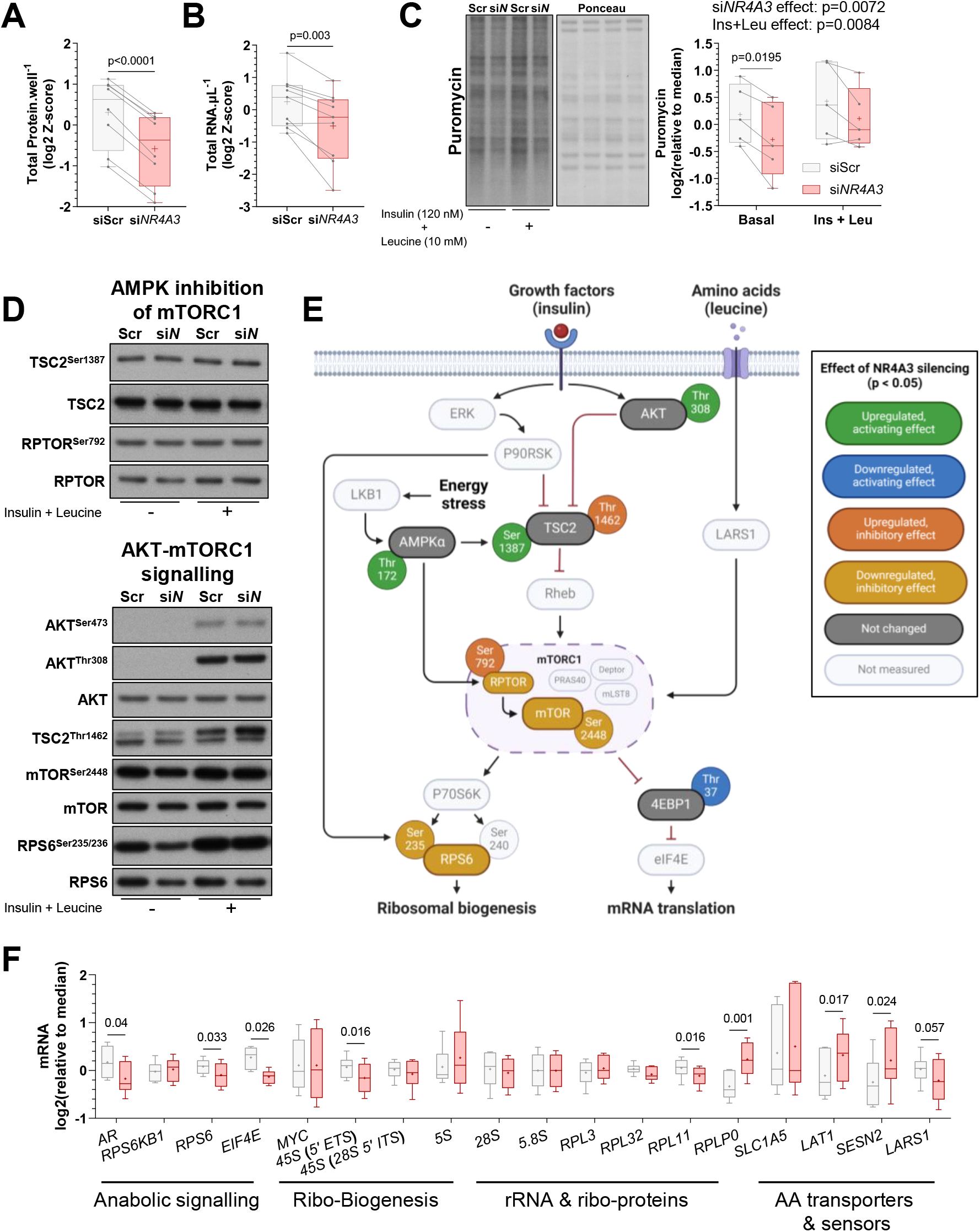
*NR4A3* silencing impairs mTORC1 signalling, ribosomal biogenesis, and translation. Primary skeletal muscle cells were exposed to a control scramble sequence (siScr) or a silencing RNA targeting *NR4A3* (si*NR4A3*). **A**. Total protein per well and **B**. RNA per μL. Results are the average Z-score of log2-transformed values across experiments. n = 8 for protein and n = 9 for RNA, paired t-test. **C**. Representative immunoblot and quantification of cellular protein synthesis assessed by protein puromycilation (i.e. SUnSET method). n = 5, 2-way ANOVA (silencing x insulin + leucine) with Šidák correction. Overall statistical model effects are stated in the figure. **D**. Immunoblot of proteins involved in AMPK-mediated mTORC1 inhibition and AKT-mTORC1 pathways from a representative donor. **E**. Schematic representation of immunoblot analysis. n = 5, 2-way ANOVA (silencing x insulin + leucine) with Šidák correction. Quantification of each depicted signalling event can be found in **Supplementary Figure 2. F**. mRNA expression of genes involved in amino acid (AA) metabolism and protein synthesis measured by RT-qPCR. Results are box-and-whisker plots with Tukey distribution and crosses indicating mean values. n = 6, paired t-tests with FDR correction. ‘Ribo-’ = ribosomal.

The efficiency and total capacity for translation in skeletal muscle is determined by mTORC1 signalling and ribosomal content, respectively (Figueiredo *et al*., 2021; West *et al*., 2016). *In vitro*, AMPK impedes mTORC1 activity through phosphorylation of tuberous sclerosis complex 2 (TSC2) (Inoki *et al*, 2003) and the regulatory-associated protein of mTOR (RPTOR) (Gwinn *et al*, 2008). Consistent with energy-stress dependent upregulation of AMPK (**Figure 3C**), the ratio of phosphorylated-to-total protein of AMPK target sites on TSC2 (Ser1387) and RPTOR (Ser792) were increased by NR4A3 depletion (**Figures 4D-E**; **Supplementary Figure 2A**) indicative of AMPK interference of mTORC1. Furthermore, *NR4A3* silencing reduced the protein abundance of RPTOR, mTOR, and ribosomal protein S6 (RPS6) (**Figures 4D-E**; **Supplementary Figures 2B-D**). These collective AMPK-related and unrelated changes coalesced to inhibit mTOR^Ser2448^ phosphorylation and activation of downstream mTORC1-substrates, including RPS6^Ser235/236^ and 4EBP1^Thr37/46^ phosphorylation, with negative consequences for skeletal muscle protein synthesis following NR4A3 depletion (**Figures 4D, 4E**; **Supplementary Figures 2C-E**).

In agreement with diminished translation, gene expression analyses confirmed that *NR4A3* RNA interference decreased mRNA abundance of *RPS6*, as well as eukaryotic translation initiation factor 4E (*EIF4E*) and 45S pre-ribosomal RNA (5’ external transcribed spacer; *5’ ETS*) (**Figure 4F**). The 45S pre-rRNA polycistronic precursor is transcribed by RNA polymerase I (Pol I) and gives rise to the 18S, 5.8S, and 28S rRNAs (von Walden, 2019). These rRNAs are subsequently assembled with Pol II-dependent ribosomal proteins and Pol III-transcribed 5S rRNA into the functional 40S (small) and 60S (large) ribosomal subunits (von Walden, 2019). As such, Pol I activity is the rate-limiting step in ribosomal biogenesis. Due to the short half-life of the ETS region (Popov *et al*, 2013), measurement of the 45S pre-RNA 5’ ETS is a reliable indicator of Pol I-mediated ribosomal transcription (von Walden, 2019). Thus, our results imply that lower levels of total RNA from *NR4A3*-silenced myotubes (**Figure 4B**) are a consequence of attenuated ribosomal biogenesis, resulting in reduced ribosomal mass, and that NR4A3 downregulation impedes both translational efficiency (i.e. mTORC1) and capacity (i.e. ribosomal abundance).

Despite an overall reduction in protein synthesis (**Figure 4C**), *NR4A3*-depleted myotubes still responded to insulin plus leucine stimulation, as suggested by the interaction between *NR4A3* silencing and treatment for mTOR^Ser2488^, RPS6^Ser235/236^, and the inhibitory AKT-target phosphorylation site on TSC2 (Thr1462) (**Figure 4D**; **Supplementary Figures 2C, 2D, 2F**). AKT^Thr308^ levels were greater upon insulin treatment (**Supplementary Figures 1A, 1E**) and tended to increase with insulin plus leucine stimulation (**Figure 4D-E**; **Supplementary Figure 2G**). Furthermore, mRNA of the large neutral amino acid transporter small subunit 1 (*LAT1*; also known as *SLC7A5*) was upregulated by *NR4A3* silencing (**Figure 4F**). LAT1 is the dominant antiporter for leucine uptake into cells (Gauthier-Coles *et al*, 2021). The rise in intracellular leucine is then detected through biochemical sensing mechanisms, including disassociation of sestrin-2 (SESN2) with GATOR2 (Saxton *et al*, 2016) and activation of leucyl-tRNA synthetase 1 (LARS1) (Han *et al*, 2012), with subsequent activation of mTORC1. As such, increased LAT1 would feasibly support higher intramuscular leucine concentrations under *NR4A3*-silenced conditions, which could have offset any detrimental effects of increased *SESN2* or reduced *LARS1* mRNA (**Figure 4F**). Hence, canonical insulin signalling, combined with enhanced leucine transport, provides a mechanism by which anabolic sensitivity of NR4A3-depleted myotubes is retained via the AKT-mTORC1 pathway.

We next investigated the impact of NR4A3 downregulation on markers of inflammation and protein degradation. Although the contribution of protein breakdown towards human disuse muscle atrophy is unclear (Brook *et al*., 2022; Pavis *et al*., 2023; Shur *et al*., 2024), the ubiquitin-proteasomal (UPS) and autophagy-lysosomal systems are markedly induced during murine models of inactivity (Min *et al*, 2011; Talbert *et al*, 2013). NR4A3 depletion upregulated activating transcription factor 3 (*ATF3*) and DNA damage-inducible transcript 3 (*DDIT3*; also known as *CHOP*) mRNA, whilst reducing IkBα protein abundance (**Supplementary Figures 3A-C**), signifying possible inflammation-mediated sarcoplasmic reticulum stress. However, other surrogates of inflammatory nuclear factor kappa B (NF-kB) signalling and the unfolded protein response, such as interleukin 6 (*IL-6*) and *ATF4*, were unaltered (**Supplementary Figure 3A**).

Major proteolytic pathways use ubiquitination as a signal for select protein degradation (Pohl & Dikic, 2019). Indeed, overexpression of the muscle-specific E3 ubiquitin ligase muscle ring-finger 1 (MuRF1 or TRIM63) is sufficient to induce ubiquitination and muscle atrophy in mice (Baehr *et al*, 2021). The *MuRF1* promoter is a direct target of forkhead box O3 (FOXO3a) (Bollinger *et al*, 2014) and inhibitory phosphorylation of FOXO3a and FOXO1 were lower after *NR4A3* interference (**Supplementary Figures 3D-E**), indicative of greater FOXO transcriptional activity. Consistent with this, mRNA expression of *MuRF1* and *FOXO3* were also elevated (**Supplementary Figure 3A**). However, these findings were contrasted by downregulation of both MURF1 and FOXO3a protein levels, while FOXO1 and the alternative muscle-enriched E3 ligase MAFbx (also known as atrogin-1) were unperturbed (**Supplementary Figures 3D-G**). Furthermore, in contrast to the stark impairment of protein synthesis, *NR4A3* silencing did not alter global protein ubiquitination in primary skeletal myotubes (**Supplementary Figure 3H**). Rather, the proteases calpain-1 (including the activated, autolysed form) and caspase 3 (inactive zymogen form) were decreased upon NR4A3 depletion, as were alpha subunit proteins of the 20S core particle proteasome (Pan 20Sα) (**Supplementary Figures 3I-K**), whereas proteasome 20S subunit alpha 1 (*PSMA1*) and proteasome 20S subunit beta 2 (*PSMB2*) transcripts were induced and attenuated, respectively (**Supplementary Figure 3A**). Additionally, despite skeletal muscle-specific *NR4A3* transgenic mice displaying greater autophagic flux (Goode *et al*, 2016), *NR4A3* interference had no impact on autophagy markers unc-51 like autophagy activating kinase 1 (ULK1), ubiquitin-binding protein p62 (also known as sequestrosome-1), or microtubule-associated proteins 1A/1B-light chain 3 (LC3-I and LC3-II) (**Supplementary Figures 3L-N**).

Altogether, silencing *NR4A3* in skeletal myotubes triggered AMPK-dependent and independent inhibition of mTORC1, reducing ribosomal biogenesis and protein translation. Pathways related to proteolysis were negligibly affected, suggesting NR4A3 downregulation primarily reduces skeletal muscle protein content via multi-level attenuation of protein synthesis.

### Overexpression of the canonical *NR4A3* isoform increases protein synthesis in primary human myotubes

The *NR4A3* gene produces four transcripts encoding three isoforms of the NR4A3 protein. The RNA interference method described in previous figures targeted an exon common to all isoforms, precluding the analysis of differential effects of select *NR4A3* variants. Thus, we explored lentiviral overexpression of the canonical *NR4A3* isoform (*NR4A3-203*; NM_006981.4) in primary human skeletal myotubes. Overexpression of the *NR4A3-203* (*NR-203*) transcript by >140-fold increased total NR4A3 protein levels by >4.5-fold without altering the abundance of other NR4A family members (**Figure 5A**) or affecting aspects of glucose and fatty acid metabolism (**Supplementary Figures 4A-D**). However, consistent with the effects of *NR4A3* silencing, *NR4A3-203* overexpression enhanced protein synthesis at baseline and after insulin plus leucine treatment (**Figure 5B**). This result was further supported by assessment of radiolabelled phenylalanine incorporation into protein (**Figure 5C**). Here, canonical *NR4A3* overexpression increased rates of translation under both unstimulated conditions and after foetal bovine serum (FBS) plus leucine treatment when combined with proteasomal blockade (lactacystin) or mTORC1-inhibition (rapamycin) (**Figure 5C**). That total cellular protein content of *NR4A3-203* overexpressing myotubes was greater in these same conditions implies the canonical NR4A3isoform may upregulate overall protein turnover in a manner partly independent from mTORC1 (**Figure 5D**). Indeed, no differences in total RNA concentrations or mTORC1 signalling were observed after *NR4A3-203* overexpression (**Supplementary Figures 4E, 4F**), suggesting that *NR4A3* silencing and overexpression phenotypes manifest through different mechanisms. This hypothesis was further substantiated by the failure of *NR4A3-203* overexpression to modulate transcripts affected by silencing (**Supplementary Figure 4G**). Nevertheless, overexpression of the canonical *NR4A3* isoform in the presence of siRNA targeting all *NR4A3* variants (**Figure 6A-C**) partially restored glucose oxidation (**Figure 6D**) concomitant with recovery of signalling downstream mTORC1: P70S6K, RPS6, and 4EBP1 (**Figure 6E, Supplementary Figures 5A-C**). Despite these effects, AMPKα^Thr172^ phosphorylation remained elevated after restoration of *NR4A3-203* (**Supplementary Figure 5D)**, which likely contributed to the sustained augmentation of fatty acid oxidation rates (**Supplementary Figure 5E**). Overall, these results posit the *NR4A3-203* isoform as a major contributor towards the negative metabolic outcomes associated with NR4A3-dependent downregulation of protein synthesis.

**Figure 5.**
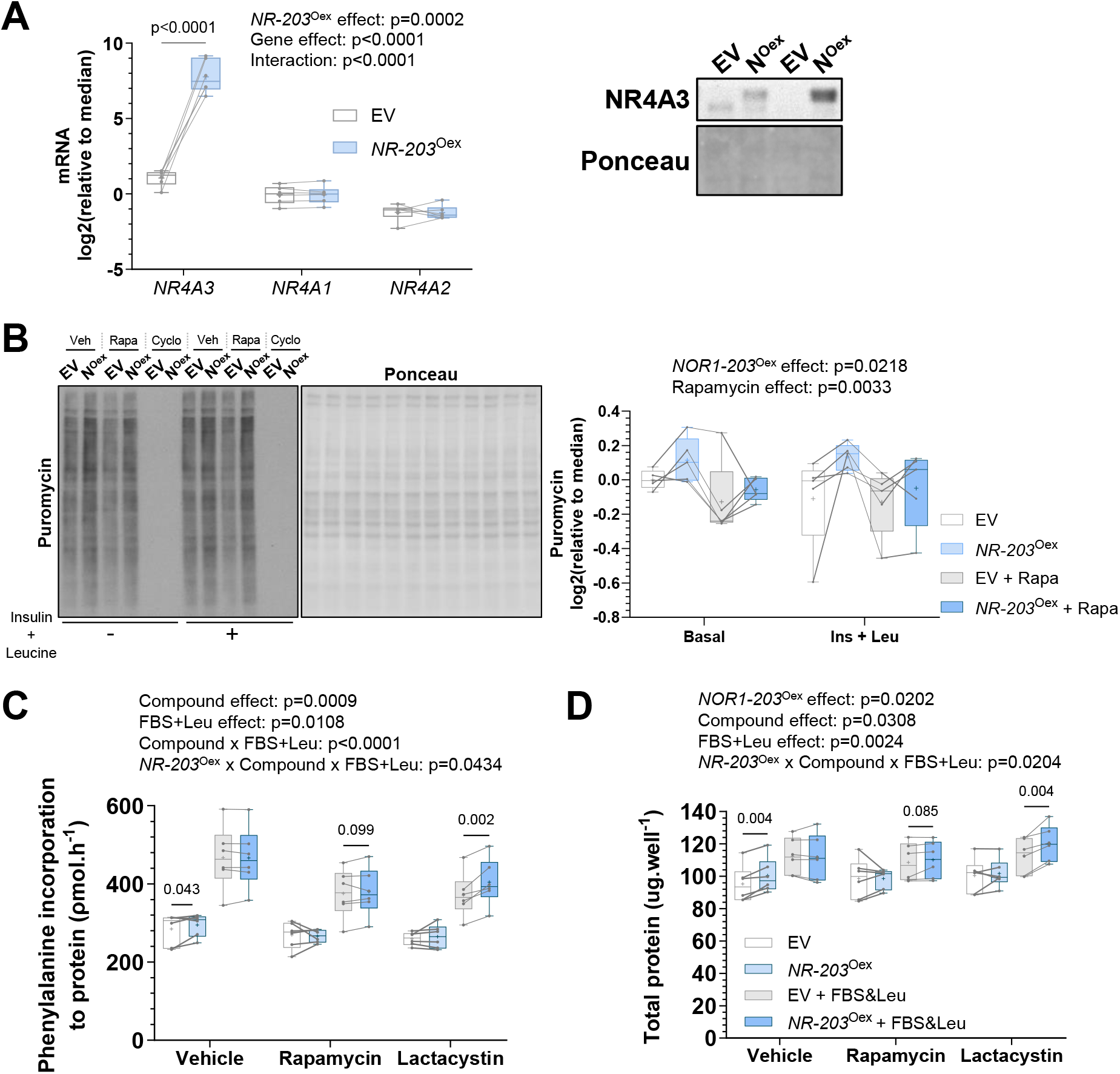
Overexpression of the canonical *NR4A3-203* isoform increases protein synthesis. Primary skeletal muscle cells were transduced with an empty vector control plasmid (EV) or a plasmid containing variant *NR4A3-203* (*NR-203*^Oex^). **A**. mRNA expression of NR4A family members *NR4A3, NR4A1* and *NR4A2* measured by RT-qPCR. n = 6, 2-way ANOVA (overexpression x gene) with Šidák correction. Also, protein level of NR4A3 from two indicative donors. **B**. Representative immunoblot and quantification of cellular protein synthesis assessed by protein puromycilation (i.e. SUnSET method). n = 5, 3-way ANOVA (overexpression x rapamycin x insulin+leucine) with FDR correction. Veh = vehicle, Rapa = rapamycin, and Cyclo = cycloheximide treatments, respectively. **C**. Rates of radiolabelled phenylalanine incorporation to protein under basal and (20%) FBS plus (10 mM) leucine-stimulated conditions over 6 h, with or without mTORC1-(rapamycin) or proteasomal-(lactacystin) inhibition. n = 6, 3-way ANOVA (overexpression x FBS+Leucine x compound) with FDR correction. **D**. Total protein per well from same experiment as C. Overall statistical model effects are stated in figures.

**Figure 6.**
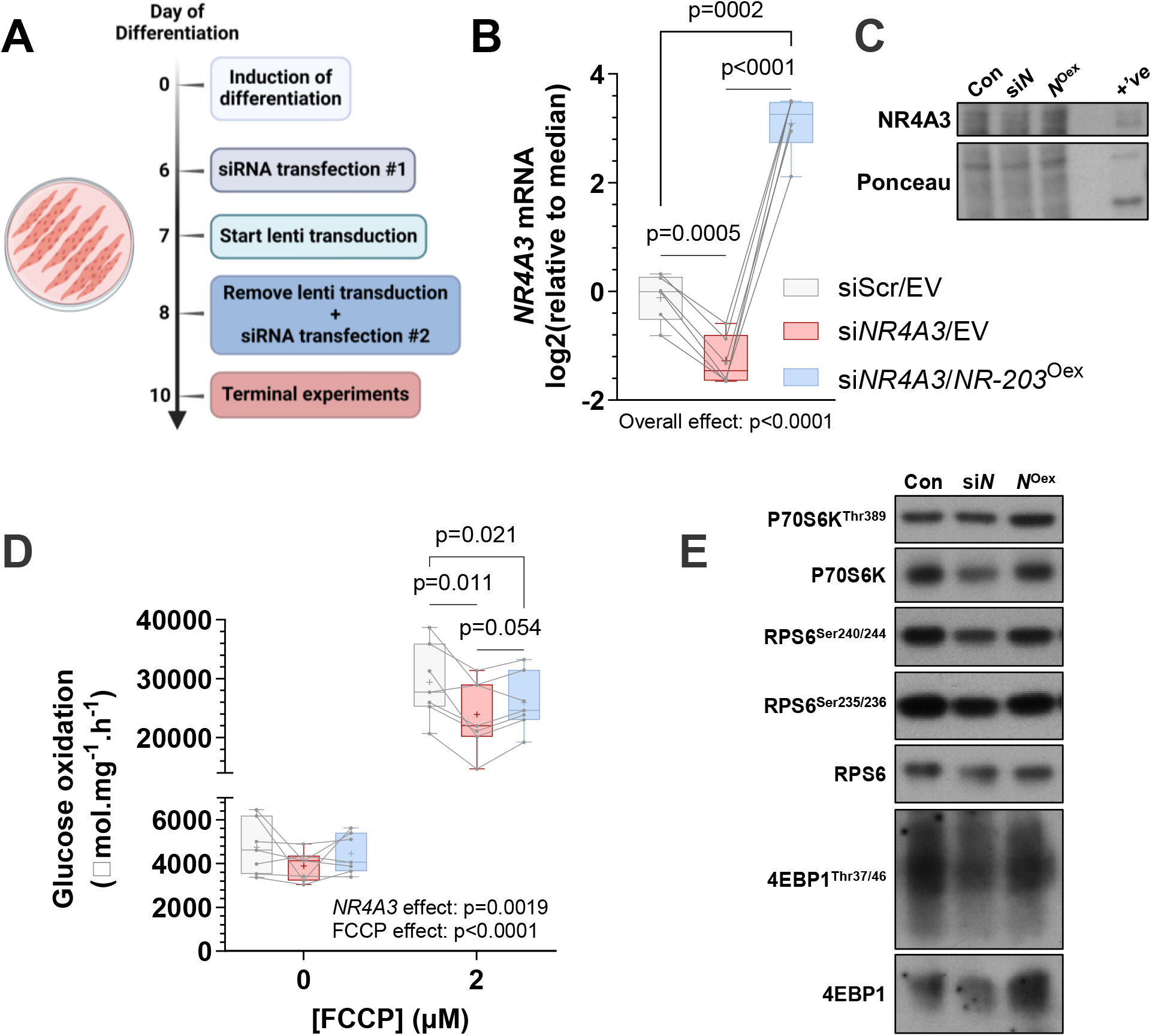
Overexpression of the canonical NR4A3 isoform partially restores glucose oxidation after global *NR4A3* silencing in myotubes, concomitant with rescued mTORC1 signalling. **A**. Schematic representation of the *NR4A3* ‘rescue’ experiment. **B**. *NR4A3* mRNA expression measured by RT-qPCR. n= 6, 1-way ANOVA with Tukey correction. **C**. Protein level of NR4A3 from one indicative donor. Con = siScr/EV, si*N* = si*NR4A3*/EV, and *N*^Oex^ = si*NR4A3*/*NR-203*^Oex^ conditions, respectively. +’ve = positive control from nuclear fraction lysate of *NR-203*^Oex^. **D**. Rates of radiolabelled glucose oxidation under basal and (2 µM) FCCP-stimulated conditions over 4 h. n = 7, 2-way ANOVA (overexpression x FCCP) with Fisher’s LSD post-test. Overall statistical model effects are stated in relevant figures. **E**. Immunoblot of proteins in the mTORC1 pathway from a representative donor. Quantification of each depicted signalling event can be found in **Supplementary Figure 5**.

### *NR4A3* silencing alters primary human myotube size and structure

*NR4A3* is decreased during inactivity associated with muscle disuse (**Figure 1A, 1B**). We therefore examined how changes in translation and protein synthesis induced by *NR4A3* silencing impact on myotube size and contractile apparatus. Immunostaining for fast-type myosin heavy chain isoforms (MYH1/2; type-IIX/type-IIA) revealed a striking decrease in myotube size (**Figure 7A**). This difference was driven by a reduction in myotube area (**Figure 7B**) without changes in the number of nuclei (**Figure 7C**) or the ability of myoblasts to fuse into myotubes (**Figure 7D**).

**Figure 7.**
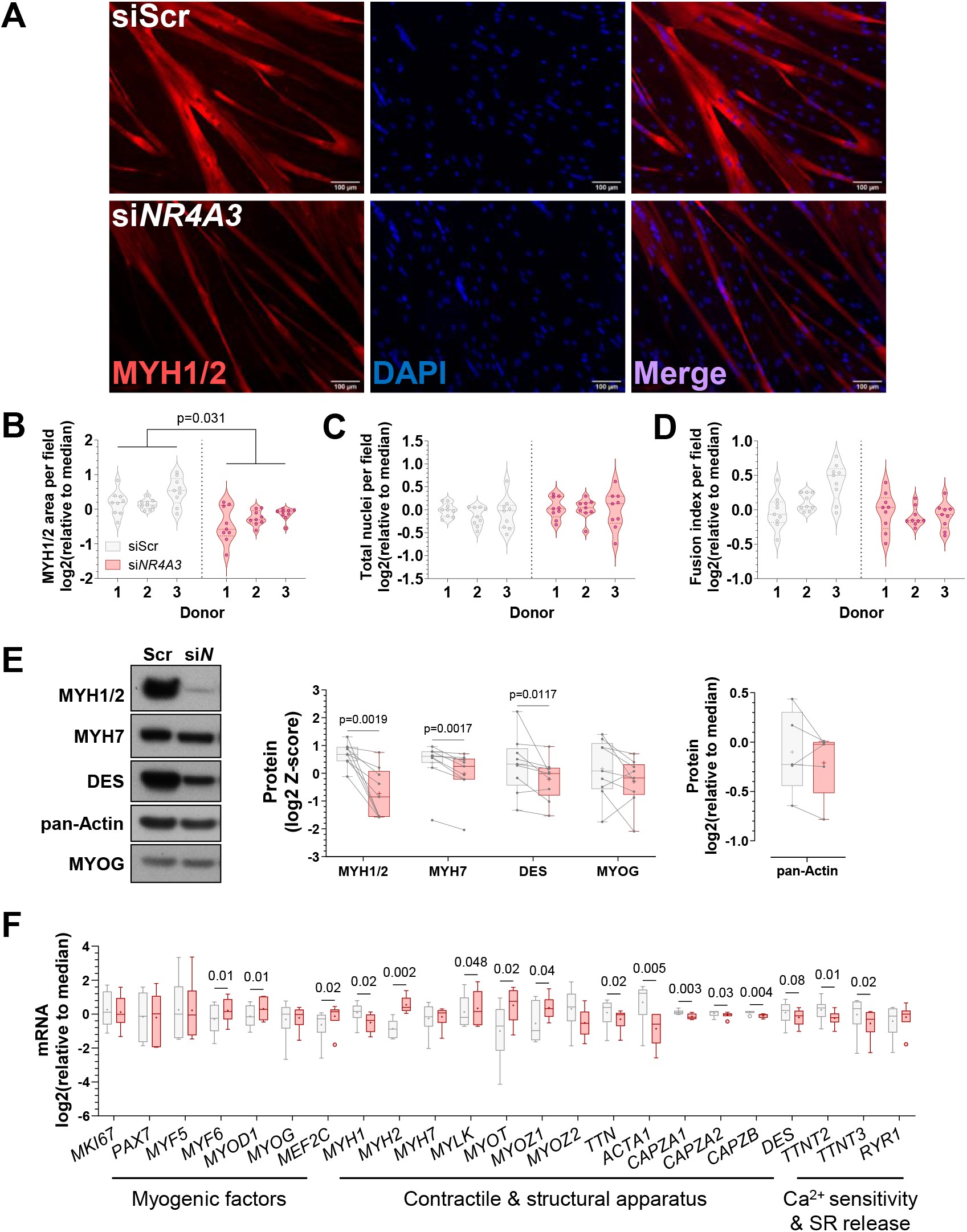
*NR4A3* silencing reduces myotube size and alters the expression of muscle contractile and structural apparatus. Primary skeletal muscle cells were exposed to a control scramble sequence (siScr) or a silencing RNA targeting *NR4A3* (si*NR4A3*). **A**. Immunocytochemistry of fast myosin heavy chain isoforms (MYH1/2, red) and nuclei (DAPI, blue) from a representative donor. Scale bar = 100 μm. Quantification of immunocytochemistry for **B**. MYH1/2 area, **C**. total nuclei, and **D**. fusion index per field of view. Results are violin plots with median and interquartile range. Circles represent measurements from different fields of view across three technical replicates per donor. n = 3, nested paired t-test. **E**. Representative immunoblot and quantification of proteins involved in contraction, structure, and myogenic transcription. Results for MYH1/2, MYH7, DES, and MYOG are the average Z-score of log2-transformed values across experiments (n = 9, paired t-test). Results for pan-Actin are n = 5, 2-way ANOVA (silencing x insulin + leucine) with only basal data shown. **F**. mRNA expression of genes involved in myogenesis, muscle structure and contraction measured by RT-qPCR. Results are box-and-whisker plots with Tukey distribution and crosses indicating mean values. n = 6, paired t-tests with FDR correction. ‘SR’ = sarcoplasmic reticulum.

NR4A3 depletion reduced the protein abundance of fast (MYH1/2) and slow (MYH7) myosin heavy chain isoforms (**Figure 7E**; **Supplemental Figure 6**) and altered the expression of transcripts encoding for proteins integral to skeletal muscle contraction and structure. This included downregulation of *MYH1*, titin (*TTN*), capping actin proteins of muscle Z-line (*CAPZ* family), and troponin isoforms *TTNT2* and *TTNT3* (**Figure 6F**). Altogether, the protein and mRNA levels of major structural and contractile proteins of skeletal muscle cells were disrupted by *NR4A3* silencing, associated with diminished myotube size.

## Conclusions

*NR4A3* is markedly induced following a single exercise bout (Pattamaprapanont *et al*., 2016), but not after habitual exercise training (Pillon *et al*., 2020). Similarly, *NR4A3* is transiently downregulated during reduced physical activity and quickly upregulated upon reloading, but unchanged after prolonged immobilisation (>2 weeks). The onset of muscle fibre atrophy occurs rapidly with disuse (Brook *et al*., 2022; Shur *et al*., 2024) and is most aggressive across the initial two weeks of unloading (Hardy *et al*, 2022), which coincides with the lowest levels of *NR4A3* mRNA. This expression pattern suggests an important role for NR4A3 in regulating skeletal muscle function during the acute transition phases associated with intense tissue remodelling. The first two weeks of unloading correspond to lower postprandial (Breen *et al*., 2013) and total daily (Shad *et al*., 2019) rates of myofibrillar protein synthesis. Here, we provide evidence that NR4A3 exerts its effects by directly influencing translation in skeletal muscle. Furthermore, NR4A3 downregulation in myotubes recapitulated adverse effects observed in human (Eggelbusch *et al*, 2024) and rodent (Siripoksup *et al*, 2024) skeletal muscle tissue following inactivity. This includes impaired glucose metabolism, reduced protein synthesis, altered expression of the contractile apparatus, and diminished myotube size. Thus, our results shed light on the involvement of NR4A3 in the control of skeletal muscle protein synthesis and metabolism during disuse atrophy.

Several lines of evidence from both *in vitro* and *in vivo* studies suggest that NR4A3 is associated with metabolic responses in skeletal muscle. Rats bred for fitness have skeletal muscle enrichment of NR4A3, associated with higher mitochondrial content and increased running capacity (Stephenson *et al*, 2012). In transgenic mice, *Nr4a3* overexpression promotes skeletal muscle remodelling towards an oxidative phenotype (Pearen *et al*., 2012), suggesting that NR4A3 plays a causal role in metabolic reprogramming of skeletal muscle. Observations from *in vivo* studies corroborate results from *in vitro* experiments in mouse C2C12 and primary human myotubes, demonstrating that silencing of *NR4A3* changes gene expression profiles towards anaerobic programmes (Pearen *et al*., 2008) and impairs mitochondrial function (Paez *et al*., 2023; Pearen *et al*., 2008; Pillon *et al*., 2020). Such effects may account for the lower ATP concentrations in C2C12 myotubes after *Nr4a3* silencing (Pearen *et al*., 2008). Our data add mechanistic evidence that NR4A3 downregulation profoundly impacts substrate utilisation, shifting metabolism towards increased lactate production and fatty acid oxidation, while decreasing glucose oxidation. Additionally, we corroborate findings in C2C12 (Paez *et al*., 2023) that the mTORC1 complex is a signalling node inhibited by NR4A3 depletion. Activation of AMPK led to the phosphorylation of TSC2 and RPTOR at sites that blunt mTORC1 transduction. However, we also observed decreased mTOR and RPS6 protein abundance, a phenomenon unlikely triggered exclusively by the activation of AMPK. Indeed, rates of translation and ribosomal biogenesis were attenuated in response to *NR4A3* RNA interference, indicating that reductions in NR4A3 impair cellular metabolism by obstructing cell-wide anabolic pathways. In agreement, rescuing protein synthesis via overexpression of the canonical NR4A3 isoform restored glucose oxidation in *NR4A3*-silenced myotubes.

Our study connects lower levels of NR4A3 with inactivity paradigms and indicates that attenuation of NR4A3 in human skeletal muscle has adverse effects on protein synthesis and metabolic responses. Muscle mass and strength are important predictors of mortality in intensive care (Weijs *et al*, 2014). Therefore, our results indicate that NR4A3 is a key molecular transducer, downregulation of which contributes to the deleterious health consequences associated with sedentary lifestyles. Understanding and addressing NR4A3 regulation could have far-reaching implications for global health and constitutes an important target to mitigate the adverse effects of inactivity, sarcopenia, and metabolic disease.

## Acknowledgments

This study was supported by Swedish Research Council (2015-00165, 2018-02389, 2023-03015),Novo Nordisk Foundation (NNF22OC0077741), Swedish Diabetes Foundation (DIA2023-824), Swedish Research Council for Sport Science (P2023-0093), Knut and Alice Wallenberg (2021.0249), Strategic Research Program in Diabetes at Karolinska Institutet (2009-1068), European Foundation for the Study of Diabetes/Novo Nordisk Foundation Future Leaders Award (NNF21SA0072747), and Diabetes Wellness Sverige (PG21-6524)

The computations were enabled by resources provided by the National Academic Infrastructure for Supercomputing in Sweden (NAISS), partially funded by the Swedish Research Council through grant agreement no. 2022-06725. Figure 4E was created with BioRender.com.

## Conflict of interest

VML is co-founder, CEO and shareholder of HepaPredict AB, as well as co-founder and shareholder of Shanghai Hepo Biotechnology Ltd. The other authors declare that they have no conflict of interest.

## Methods

### Transcriptomic meta-analysis

Transcriptomic studies of inactivity in human trials were selected using the MetaMEx database (Pillon *et al*., 2020). Studies were annotated with the sex and age of participants, as well as the total duration of inactivity. Statistics were performed individually for each study and the mean, variance and *n* size were used to fit a random effects model to the data. The restricted maximum-likelihood (REML) method was applied using the R package metaphor. Obtained p-values were adjusted using the Benjamini & Hochberg method. To evaluate the effect of inactivity duration, all transcriptomic studies were merged by gene name and split in three groups: less than one week, one-two weeks or more than two weeks. A linear model was used to test the overall effect of inactivity duration, as well as pair-wise comparisons to the pre-inactivity group, using Empirical Bayes Statistics for Differential Expression with the limma package (Ritchie *et al*, 2015). Gene set enrichment analysis was performed using ClusterProfiler (Wu *et al*, 2021) on genes ranked based on Spearman correlation with *NR4A3*. Gene ontology biological processes were considered significant at FDR<0.05. The effect of reloading was estimated with the same method as above after merging GSE21496 and GSE24215.

### Primary human cell culture

Primary cells were isolated from *vastus lateralis* skeletal muscle biopsies derived from ten healthy volunteers (age: 36 ± 15 years; BMI: 24.0 ± 1.9 Kg.m-^2^; biological sex: eight males, two females). The regional ethical review board in Stockholm approved protocols. Cells were cultured and differentiated as described previously (Abdelmoez *et al*., 2020). Successful myotube formation was monitored under the microscope and cells were used for terminal experiments ten days after the initiation of differentiation. The absence of mycoplasma contamination was routinely confirmed by PCR.

### Silencing by RNA interference

On days six and eight of differentiation, myotubes were transfected with 10 nM of either Silencer Select siRNA Negative Control No. 2 (no. 4390847) or Silencer Select siRNA s15542 targeting *NR4A3* (Life Technologies, Foster City, CA). Transfections were performed for 5 h in OptiMEM reduced serum media with Lipofectamine RNAiMAX transfection reagent (Invitrogen, Carlsbad, CA). Terminal experiments were performed ≈48 h after the second transfection.

### Plasmid transformation, expansion, and purification

cDNA of the canonical *NR4A3* transcript (NR4A3-203; NM_006981.4) was synthesised into a modified pLenti CMV Puro DEST (w118-1) vector (Addgene plasmid #17452) by GENEWIZ (Azenta Life Science, Leipzig, Germany). One Shot TOP10 Chemically Competent *E. coli* cells (Invitrogen, ThermoFisher Scientific) were used for bacterial transformation and expansion. Modified pLenti CMV Puro DEST (w118-1) empty vector (EV) and NR4A3-overexpressing (*NR-203*^Oex^) plasmid DNA was then purified using endotoxin-free buffers (Cat. No. 19048) and columns (Cat. No. 10083) from Qiagen, before assessing DNA concentration and purity by spectrophotometry.

### Lentivirus production

HEK293T cells were grown in DMEM media (Gibco #10569) supplemented with 10% foetal bovine serum, 1% penicillin-streptomycin, and 500 μg/mL geneticin (G418). Twenty-four hours prior to transfection, HEK293T cells were seeded into T-225 flasks coated with poly-L-lysine. Once ≈90% confluent, lentiviral harvest was conducted by transfecting each T-225 with a transfection reagent mix consisting of 69 μg of EV or *NR-203*O^ex^ plasmid DNA, second generation packaging vectors (69 μg of psPAX2 DNA and 45 μg of pMD2.G DNA), 549 μg polyethylenimine (PEI; 3 μg per μg of total plasmid DNA) and PBS at a final volume of 4.5 mL. The transfection mix was then added to 40.5 mL of pre-warmed growth medium (without geneticin) and added to HEK293T cells for 12-14 h. At this point, the transfection media was removed and replaced with fresh medium for 2 h before switching to new growth media supplemented with 5 mM sodium butyrate. Thirty hours later, viral media was collected and passed through a 0.45 μM low-protein binding filter (Millipore) and concentrated by low-speed centrifugation (3300 g for 30 min at 4°C) using 100 kDa molecular weight cut-off Centricon Plus-70 cartridges (UFC710008). Three volumes of concentrated virus medium were subsequently mixed with one volume of Lenti-X concentrator (Takara Bio) and rotated at 4°C overnight before centrifugation (1500 g for 45 min at 4°C). Supernatant was discarded and the pellet resuspended in PBS, aliquoted, and stored at - 80°C. Virus concentration was determined using Lenti-X GoStix Plus (Takara Bio) after one freeze-thaw cycle to be consistent with experimental conditions.

### Transduction of primary skeletal myotubes

On day 7 of differentiation, myotubes were transduced with either EV or *NR-203*^Oex^ lentivirus (concentration equivalent to 16.5 ng of p24.mL-^1^ per 9.6 cm^2^ of cells) in post-fusion medium supplemented with 5 ug.mL^-1^ of polybrene. After 18-20 h, viral medium was removed, myotubes were washed once with PBS and then switched to normal post-fusion media (Abdelmoez *et al*., 2020). Terminal experiments were performed ≈48 h later.

### RNA extraction and analysis

RNA was isolated from cultured muscle cells using TRIzol-cholorform extraction according to manufacturer’s instruction (Invitrogen). RNA concentration and purity were determined by spectrophotometry. All equipment, software, and reagents for performing reverse transcription and RT-qPCR were from Thermo Fisher Scientific. The High-Capacity cDNA Reverse Transcription kit was used for cDNA synthesis according to manufacturer’s instructions. RT-qPCR was performed on a StepOne Plus system using Taqman or Sybr Green technologies (probe sequences for all analysed genes are available upon request). Relative gene expression was calculated by the comparative ΔΔCt method using the geometric mean of at least two of the following reference genes: *18S, B2M, GUSB, HPRT1* or *TBP*. The most stable reference gene combination was determined within experimental conditions.

### Glucose uptake

Myotubes were washed once with sterile PBS and incubated for 4 h in unsupplemented, serum-free low-glucose Dulbecco’s Modified Eagle Medium (DMEM) (5.5 mM glucose, Gibco #21885) (i.e. serum starvation medium). Cells were then switched to fresh serum starvation medium in the absence or presence of insulin (120 nM) for 1 h. Myotubes were washed once with warm glucose- and serum-free DMEM (Gibco #11966) and glucose uptake was measured in the same medium by adding 1 uL.mL^-1^ of 2-[1,2-^3^H]Deoxy-D-glucose (MT911, Moravek) and 10 μM unlabelled 2-Deoxy-D-glucose for 15 min (Abdelmoez *et al*., 2020). Cell monolayers were washed three times with ice-cold PBS and lysed in 1 mL 0.03% SDS. 0.5 mL of the cell lysate was counted in a liquid scintillation counter (TRI-CARB 4910TR, PerkinElmer) and protein content of the remaining lysate was measured for normalisation of results (BCA Protein Assay Kit; #23225, Thermo Fisher Scientific, Rockford, IL).

### Glycogen synthesis

Incorporation of glucose into glycogen was performed using similar methods detailed elsewhere (Abdelmoez *et al*., 2020). Briefly, myotubes were nutrient-starved as in glucose uptake experiments and then treated with 0, 10, or 120 nM of insulin in fresh serum-starvation medium for 30 min before adding 2 μL.mL^-1^ D-[^14^C(U)] glucose (NEC042B005MC; Perkin Elmer) for a further 90 min. Cells were washed three times with ice-cold PBS and lysed in 0.5 mL 0.03% SDS. Thereafter, [^14^C]-labelled glycogen was extracted from 0.4 mL of the lysate and counted in a liquid scintillation counter (Abdelmoez *et al*., 2020). Cellular protein concentration was determined by bicinchoninic acid (BCA) assay from the remaining 0.1 mL of cell lysate and used to normalize results.

### Glucose oxidation

Glucose oxidation was performed using methods as described (Abdelmoez *et al*., 2020). Myotubes were washed with PBS and incubated with D-[^14^C(U)] glucose in serum starvation medium in the presence or absence of 2 μM carbonyl cyanide p-trifluoromethoxyphenylhydrazone (FCCP) for 4 h. Thereafter, [^14^C]-CO_2_ was released from the medium by adding of 150 μL 2 M HCl per mL of medium and captured in 300 μL of 2 M NaOH for 1h. The captured [^14^C]-CO_2_ was counted using a liquid scintillation counter. Cells were washed three times with ice-cold PBS and homogenised in 0.4 mL of 0.5 M NaOH. After complete lysis, 0.1 mL of 2 M HCl was added to neutralise pH and protein was assessed using the BCA assay for normalisation of results.

### Lactate assay

Lactate released into culture supernatant was measured in post-fusion medium (Abdelmoez *et al*., 2020) after 48 h of basal or (2 μM) FCCP-stimulated conditions using a colourimetric assay as described (Abdelmoez *et al*., 2020). Assay buffer contained Tris (50 mM pH 8), NAD (7.5 mM), N-methylphenazonium methyl sulfate (250 µM), p-iodonitrotetrazolium violet (500 µM), and lactate dehydrogenase (4 U.mL^-1^). Medium was passed through 3 kDa filters by centrifugation (14000 g for 15 min at 4°C) and 50 μL of sample was mixed with 150 µL of assay buffer. After a 20-min incubation the absorbance was read at 490 nm. Results were normalised to total cellular protein content as measured by BCA assay after lysis in 0.03% SDS.

### Fatty acid oxidation

[^14^C(U)] palmitic acid (NEC534050UC; Perkin Elmer) oxidation was performed using the same experimental procedure as glucose oxidation, except that 1 μL.mL^-1^ of isotope was added to base glucose-free DMEM (Gibco #11966) supplemented with 25 μM of unlabelled BSA-conjugated palmitate.

### Thin-layer chromatography (TLC)

Myotubes were incubated for 6 h in the same medium as described for [^14^C(U)] palmitic acid oxidation. Cells were then washed three times with ice-cold PBS and hydrophobic lipids were extracted as detailed elsewhere (Massart *et al*, 2014). The resulting lipid pellet was eluted in 50 μL of a methanol:chloroform solution (1:1) and loaded on a TLC plate (Silica Gel G 250 μm 20×20 cm, Analtech, DE, USA). Lipid species were separated for 30 min in a loading chamber filled with 100 mL of a hexane:diethylether:acetic acid mixture (80:20:3). The loaded TLC plate was then transferred to an exposure cassette (GE Healthcare) and exposed to an X-ray film for four weeks at -80°C before developing in an X-ray developer machine. Identification of the appropriate bands was determined against standards for 1,2-Dioctanyl [1-^14^C] rac-glycerol (1,2-DAG), 1,3-Dioleoyl-rac-glycerol [oleoyl-1-^14^C] (1,3-DAG) (American Radiolabeled Chemicals Inc.), [^14^C(U)] palmitic acid (Perkin Elmer), and [^14^C] triolein (TG). Densitometric quantification was performed using Image Lab software (Bio-Rad).

### Phenylalanine incorporation into protein

Myotubes were washed once with PBS and incubated in low-glucose base DMEM for 4 h. After starvation, media was changed to low-glucose DMEM supplemented with an additional 25 μM unlabelled phenylalanine and 4 μL of [^14^C]phenylalanine (NEC284E050UC; Perkin Elmer) per mL for 6 h, with or without 20% FBS plus 10 mM leucine, and in the presence or absence of 100 nM rapamycin or 10 μM lactacystin. Cells were then washed three times with ice-cold PBS and lysed in 0.5 mL 0.03% SDS. Protein concentration was measured from 0.1 mL of cell lysate by BCA protein assay. Protein from the remaining cell lysate was precipitated in 50% Trichloroacetic acid with 1% BSA overnight at −20°C, followed by centrifugation (12000 g for 15 min at 4°C). Supernatant was discarded and the protein pellet washed twice in acetone with centrifugation after each wash (12000 g for 15 min at 4°C). Acetone was discarded and the protein pellet dissolved in 0.5 M NaOH for 1 h in a heating block at 65°C with shaking. The dissolved pellet homogenate was transferred to scintillation vials and the amount of [^14^C] phenylalanine incorporation determined by scintillation counting.

### Subcellular fractionation

Separation of sarcoplasmic and nuclear fractions was performed on fresh cells, without freezing. Myotubes were washed once in ice-cold PBS and scraped in cell lysis buffer (10 mM HEPES pH 7.5, 10 mM KCl, 0.1 mM EDTA, 1 mM DTT, 0.5% NP-40) supplemented with 1 mM PMSF, 0.5 mM Na_3_VO_4_ and 1x Protease Inhibitor Cocktail Set 1 (PIC) (Calbiochem). Lysates were mixed by pipetting and rotated at 4°C for 20 min then subjected to centrifugation (12000 g for 10 min at 4°C). The sarcoplasmic fraction (supernatant) was collected and stored at -80°C. The remaining pellet was washed three times in ice-cold cell lysis buffer by inverting the Eppendorf multiple times per wash and then resuspended in nuclear extraction buffer (20 mM HEPES pH 7.5, 400 mM NaCl, 1 mM EDTA, 1 mM DTT) supplemented with 1 mM PMSF, 0.5 mM Na_3_VO_4_ and 1x PIC. Resuspension was performed in the same volume collected for the sarcoplasmic fraction to retain the sarcoplasmic to nuclear ratio of proteins. Samples were next rotated at 4°C for 30 min and debris pelleted by centrifugation (12000 g for 15 min at 4°C). The nuclear fraction supernatant was collected and stored at -80°C until lysate preparation for immunoblotting.

### Surface sensing of translation (SUnSET) assay

Relative rates of in vitro protein synthesis were determined using the SUnSET method as described (Goodman *et al*., 2011). Briefly, myotubes were washed once in PBS and incubated in low-glucose base DMEM for 4 h. Media was then changed to fresh low-glucose vehicle DMEM or low-glucose DMEM supplemented with 120 nM insulin and 10 mM leucine, with or without 100 nM rapamycin or 178 µM cycloheximide (*NR-203*^Oex^ experiment only). After 1 h of treatment, puromycin (1 μM) was added to the medium and cells were incubated for a further 30 min before collection. Medium was aspirated and cells were washed once in ice-cold PBS. Samples were then harvested in protein lysis buffer (137 mM NaCl, 2.7 mM KCl, 1 mM MgCl_2_, 1% Triton X-100, 10% glycerol, 20 mM Tris pH 7.8,10 mM NaF, 1 mM EDTA, 1 mM PMSF, 0.5 mM Na_3_VO_4_ and 1x PIC), rotated for 30 min at 4°C and subjected to centrifugation (12,000 g for 15 min at 4°C). Supernatants were stored at -80°C until protein determination by BCA assay and subsequent sample preparation for immunoblotting.

### Insulin stimulation for assessment of signalling

For interrogation of the canonical insulin signalling cascade, myotubes were incubated in serum starvation medium for 4 h. Cells were then stimulated with 0, 10, or 120 nM of insulin in fresh serum starvation medium for 20 min and lysed as mentioned for SUnSET assay samples.

### Immunoblot analysis

Samples were prepared for SDS-PAGE with Laemmli buffer (60 mM Tris pH 6.8, 2% w/v SDS, 10% v/v glycerol, 0.01% w/v bromophenol blue, 1.25% v/v β-mercaptoethanol). Equal amounts of protein were loaded and separated on Criterion XT Bis-Tris Gels (Bio-Rad, Hercules, CA) and transferred to polyvinylidene fluoride membranes (Merck, Germany). Membranes were stained with Ponceau S to confirm transfer quality and even loading of samples. Membranes were then blocked with 5% non-fat milk in Tris-buffered saline supplemented with Tween-20 (TBST; 20 mM tris-HCl pH 7.6, 137 mM NaCl, 0.02% Tween-20) for 1 h at room temperature, before an overnight incubation with primary antibodies at 4°C with gentle rocking. Primary antibodies were diluted in TBS plus 0.1% w/v bovine serum albumin and 0.1% w/v NaN_3_ or 5% non-fat milk in TBST (specific antibodies and dilutions are available upon request). Membranes were next washed with TBST and incubated with species-appropriate horseradish peroxidase-conjugated secondary antibody (1:25000 in TBST with 5% non-fat milk). Proteins were visualised by enhanced chemiluminescence (Amersham ECL Western Blotting Detection Reagent, UK) and quantified by densitometry (QuantityOne or Image Lab software, Bio-Rad).

### Total protein and RNA

Unless stated otherwise, total protein concentrations reported in figures are from BCA protein assay measurements of basal conditions in metabolic assays. Compared to sample lysis for immunoblot analysis,homogenisation of cell monolayers for normalisation of metabolic assays was performed in larger volumes of buffer. Thus, these measurements more accurately reflect total protein concentration per well. In cases where total protein and RNA was determined for multiple experiments using primary cells from the same donor, values were log2 transformed, Z-scored and the average Z-score across assays was used for statistical analysis.

### Immunocytochemistry

Myotubes were immunostained for fast myosin heavy chain isoforms (MYH1/2, 1:250; sc-53088, Santa Cruz Biotechnology) using methods as described (Abdelmoez *et al*., 2020), with the addition of nuclear counterstaining. After 90 min incubation with Alexa Fluor 594 goat anti-mouse (A-11005) secondary antibody (1:500) at room temperature with gentle rocking, cells were washed twice with 0.025% Tween-20 in PBS and counterstained with 300 nM 4′,6-diamidino-2-phenylindole (DAPI) in PBS for 5 min at room temperature. Myotubes were then washed twice in PBS and kept in PBS protected from light at 4°C until imaging. MYH1/2 and nuclei images were taken using 20x magnification on a Zeiss Axio Vert.A1 inverted fluorescent microscope, equipped with ZEN software (Carl Zeiss Microscopy GmbH, Germany). Images were obtained from three random fields of view per well across 3 technical replicate wells, producing nine images per condition for each donor. MYH1/2 images were converted to 8-bit binary and quantification of MYH1/2 area (μm^2^) per field was automated using ImageJ software (Fiji) to prevent measurement bias. Nuclei were counted manually using the ImageJ cell-counter plugin. Fused myonuclei were considered nuclei counted within the MYH1/2 area automatically detected by ImageJ software in the MYH1/2 analysis and fusion index calculated as the percentage of fused versus total nuclei per field of view.

### Statistics

Analyses were performed using either R 4.1.0 (www.r-project.org) or GraphPad Prism 10.0.3 software (GraphPad Software Inc.). Exact p-values are specified for comparisons where probability met p<0.1. Data are presented as box-and-whisker plots with Tukey distribution and pairing by donor unless specified otherwise in figure legends. Normality was assessed using Shapiro-Wilk test before applying appropriate parametric or non-parametric tests. In 2-way ANOVA analyses, Tukey ladder transformation was used when residuals violated assumptions of normality. Statistical tests and sample sizes are described in figure legends. Bioinformatic analyses are described in their respective method sections.

## Supplementary Figures

**Supplementary Figure 1.**
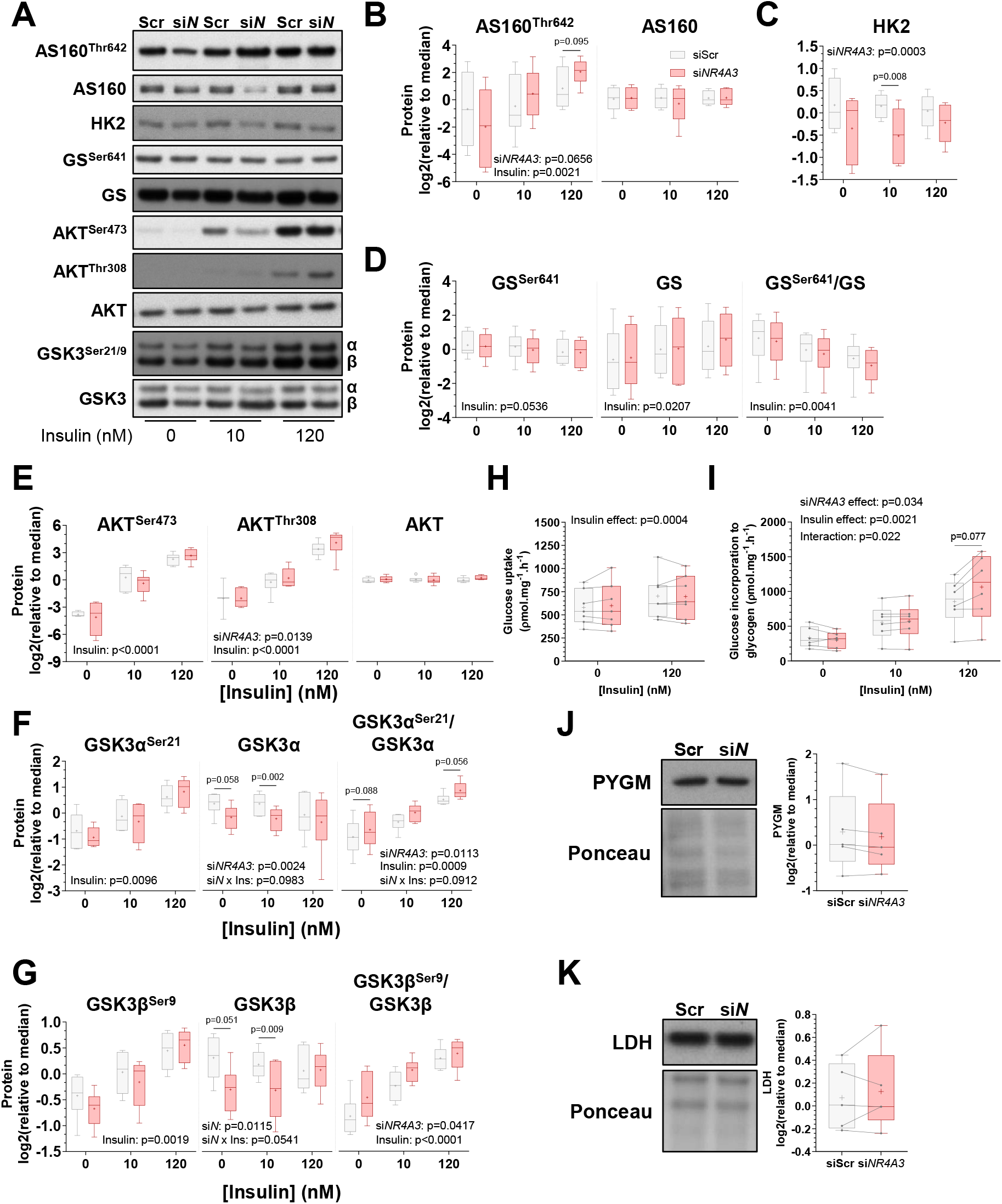
*NR4A3*-silenced myotubes retain responsiveness to insulin stimulation. Primary skeletal muscle cells were exposed to a control scramble sequence (siScr) or a silencing RNA targeting *NR4A3* (si*NR4A3*). **A**. Representative immunoblot and **B-G**. quantification of proteins and phosphorylation events in the canonical insulin signalling cascade. n = 6, 2-way ANOVA (silencing x insulin) with Šidák correction. Overall statistical model effects are stated in figures. **H**. Rates of radiolabelled 2-Deoxy-D-glucose uptake (n = 7) and **I**. glycogen synthesis (n = 6) under basal and insulin-stimulated conditions. 2-way ANOVA (silencing x insulin) with Šidák correction. **J**. Representative immunoblot and quantification of total glycogen phosphorylase, muscle associated (PYGM) and **K**. lactate dehydrogenase (LDH). n = 5, 2-way ANOVA (silencing x insulin + leucine). Only basal data are shown.

**Supplementary Figure 2.**
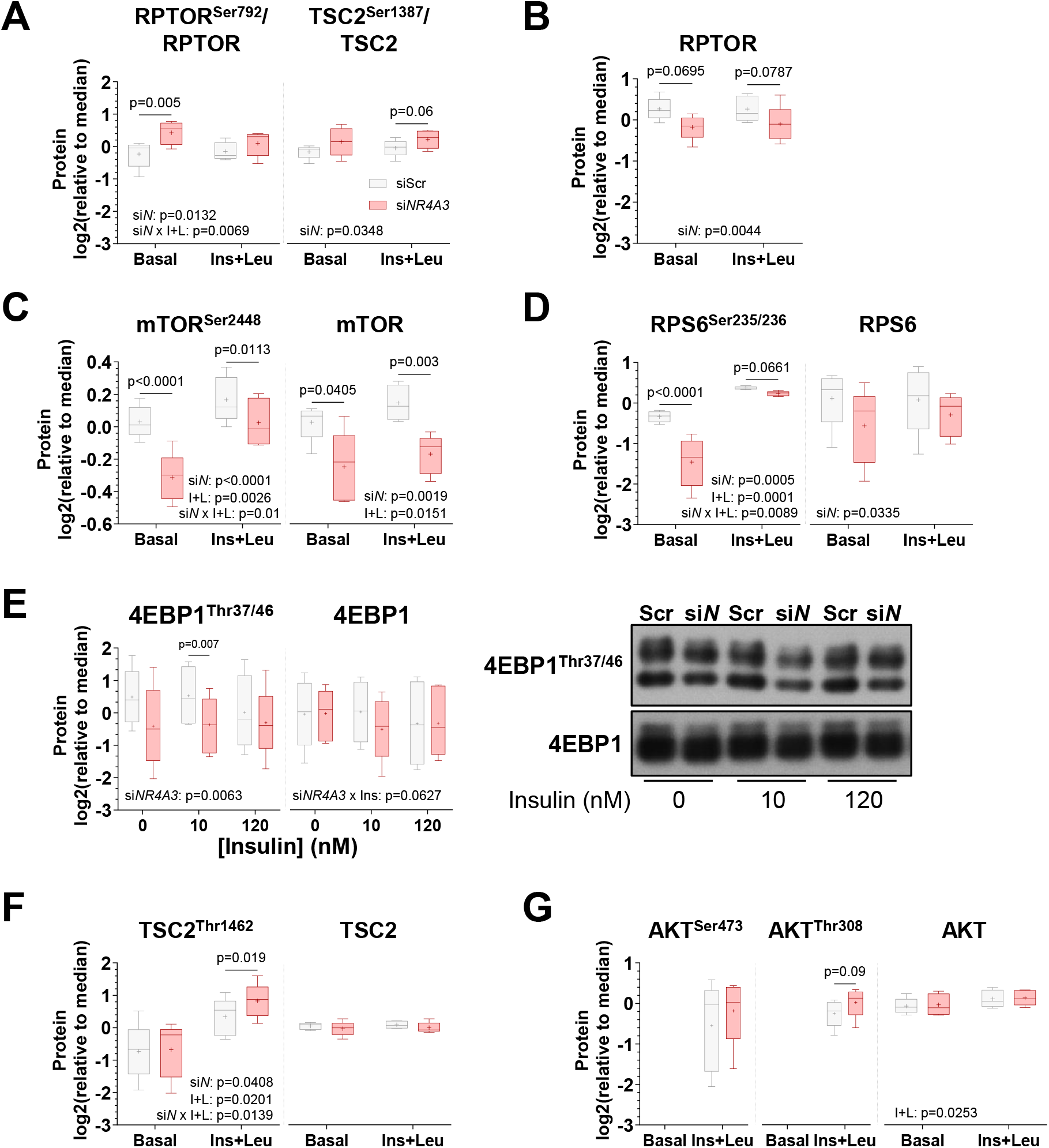
*NR4A3* silencing attenuates mTORC1 signalling. Primary skeletal muscle cells were exposed to a control scramble sequence (siScr) or a silencing RNA targeting *NR4A3* (si*NR4A3*). **A**. Immunoblot quantification of phosphorylated-to-total protein ratios for AMPK target sites on RPTOR (Ser792) and TSC2 (Ser1387). **B**. Immunoblot quantification of AKT-mTORC1 activating phosphorylation sites and total protein levels for RPTOR, **C**. mTOR, **D**. RPS6, **E**. 4EBP1 (with images from a representative donor), **F**. TSC2, and **G**. AKT. All immunoblot analyses are n = 5, 2-way ANOVA (silencing x insulin + leucine) with Šidák correction; except 4EBP1, which is n = 6, 2-way ANOVA (silencing x insulin) with Šidák correction, and AKT^Ser473^ and AKT^Thr308^ phosphorylation, which are n = 5, paired t-tests. Overall statistical model effects are stated in figures. si*N* = main silencing effect, I+L = main insulin + leucine effect, si*N* x I+L = interaction effect.

**Supplementary Figure 3.**
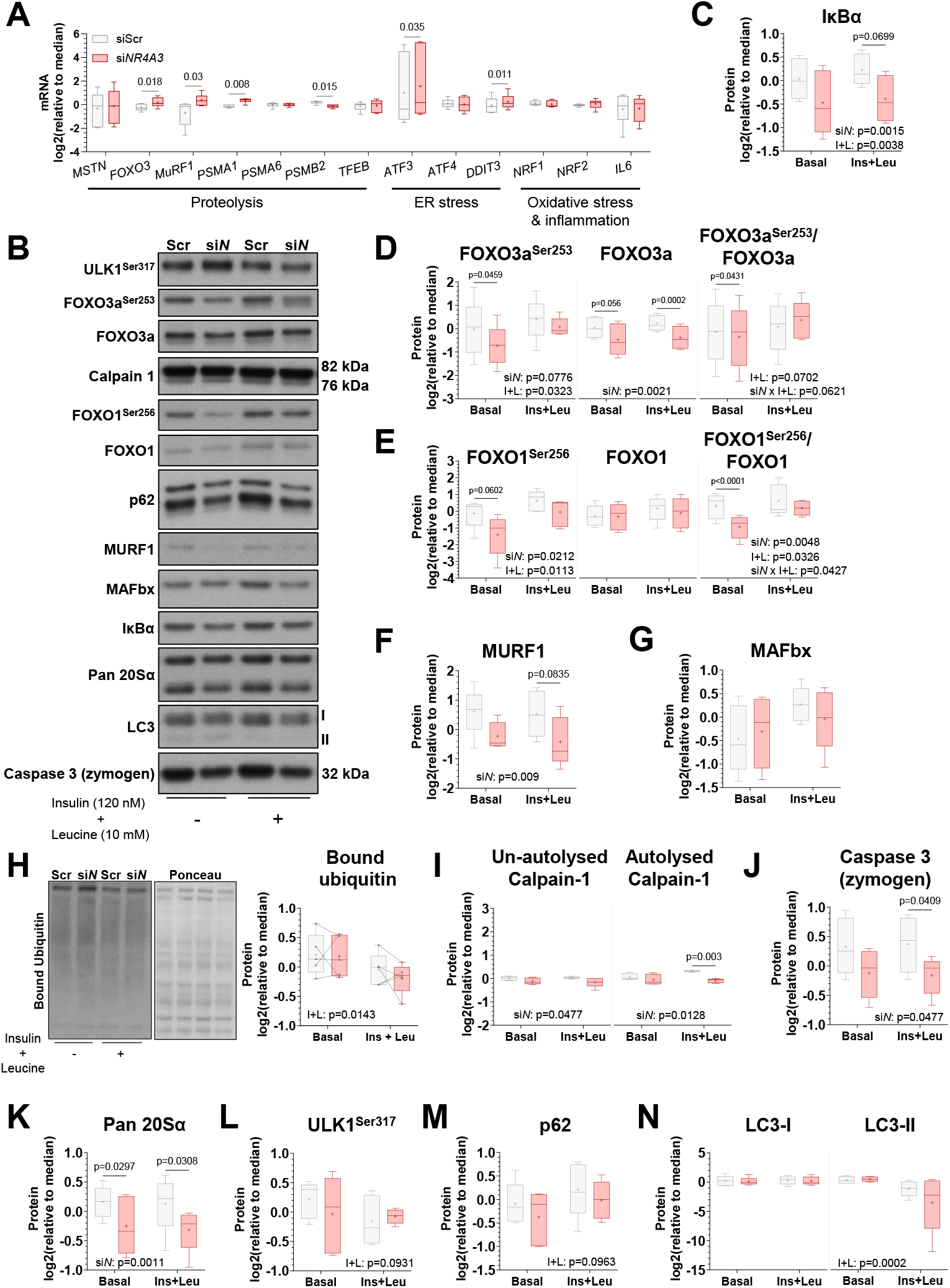
*NR4A3* silencing minimally impacts pathways controlling protein degradation. Primary skeletal muscle cells were exposed to a control scramble sequence (siScr) or a silencing RNA targeting *NR4A3* (si*NR4A3*). **A**. mRNA expression of genes related to the negative regulation of muscle mass, endoplasmic reticulum (ER) stress, oxidative stress, and inflammation measured by RT-qPCR. Results are box-and-whisker plots with Tukey distribution and crosses indicating mean values. n = 6, paired t-tests with FDR correction. **B**. Immunoblot of proteins involved in inflammation, ubiquitin-proteasomal, and autophagy-lysosomal degradation pathways from a representative donor. **C-G**. Immunoblot quantification of **C**. NF-κB inhibitor IκBα, basal and insulin-mediated inhibitory phosphorylation of transcription factors **D**. FOXO3a, **E**. FOXO1, and E3 ubiquitin ligases **F**. MURF1 and **G**. MAFbx. **H**. Representative immunoblot and quantification of cellular protein ubiquitination. **I-K**. Immunoblot quantification of **I**. Calpain-1 and **J**. Caspase 3 proteases, and **K**. alpha subunits of the 20S core particle proteasome (Pan 20Sα). **L-N**. Immunoblot quantification of autophagy markers **L**. ULK1 phosphorylation at Ser317, and total **M**. p62 and **N**. LC3-I/LC3-II abundance. All immunoblot analyses are n = 5, 2-way ANOVA (silencing x insulin + leucine) with Šidák correction. Overall statistical model effects are stated in figures.

**Supplementary Figure 4.**
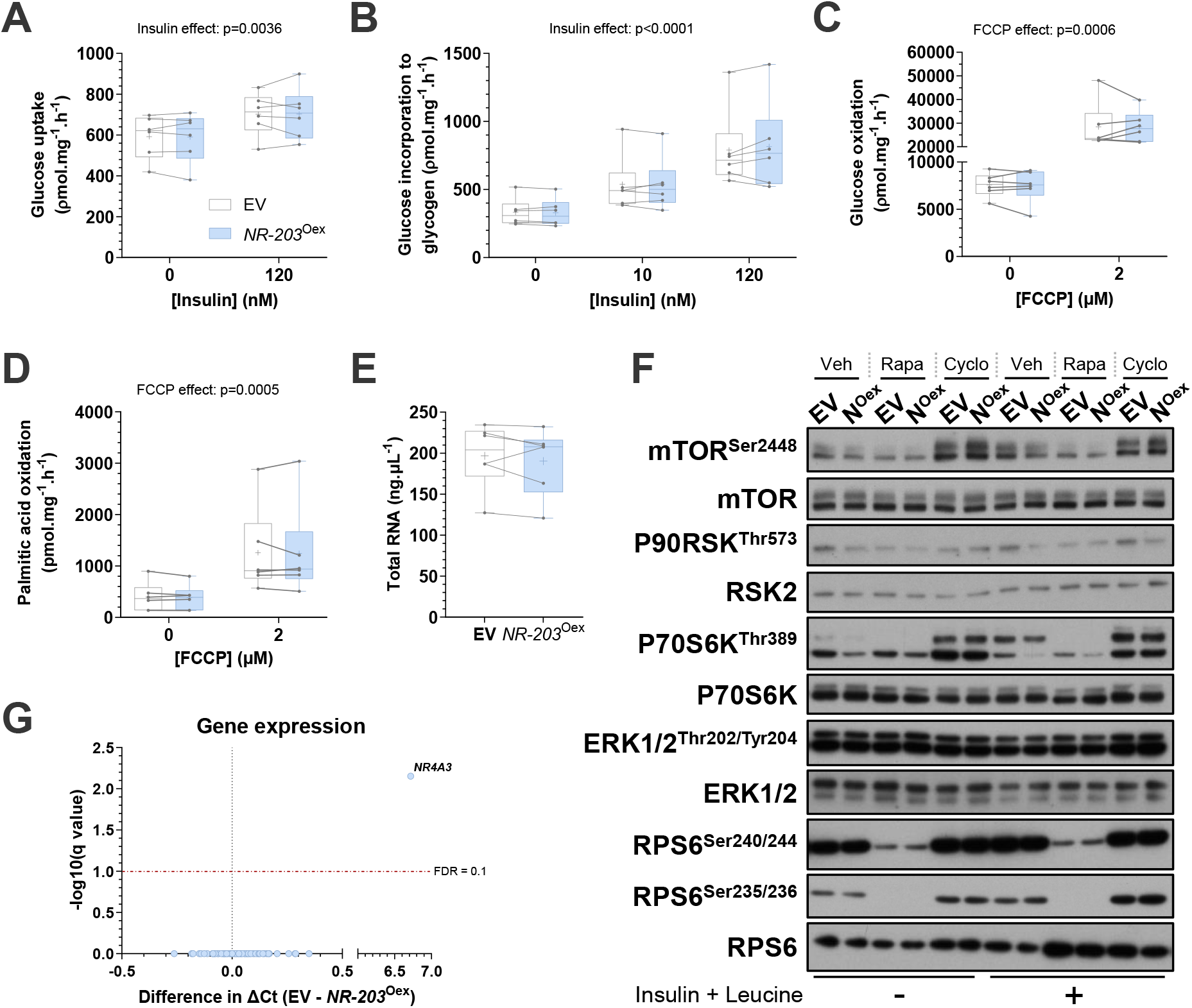
Overexpression of the canonical *NR4A3* isoform increases translation without altering glucose and fatty acid metabolism, RNA abundance, or mTORC1 signalling. Primary skeletal muscle cells were transduced with an empty vector control plasmid (EV) or a plasmid containing variant *NR4A3-203* (*NR-203*^Oex^). **A**. Basal and stimulated rates of radiolabelled 2-Deoxy-D-glucose uptake, **B**. glycogen synthesis, **C**. glucose oxidation, and **D**. ^14^C palmitic acid oxidation. n = 6, 2-way ANOVA (silencing x stimulation). Overall statistical model effects are stated in figures. **E**. Total RNA concentration (ng.μL^-1^). n = 6, paired t-test. **F**. Immunoblot of proteins in the mTORC1 and ERK signalling cascades from a representative donor. Veh = vehicle, Rapa = rapamycin, and Cyclo = cycloheximide treatments, respectively. Samples are those presented in main **Figure 5B**. No difference in phosphorylation or total protein levels were detected with *NR-203*^Oex^. n = 5, 3-way ANOVA (overexpression x rapamycin x insulin + leucine). **G**. Volcano plot of all mRNA transcripts assessed after *NR-203*^Oex^ measured by RT-qPCR. Genes were the same as analysed in *NR4A3*-silenced conditions. n = 6, paired t-tests with FDR correction.

**Supplementary Figure 5.**
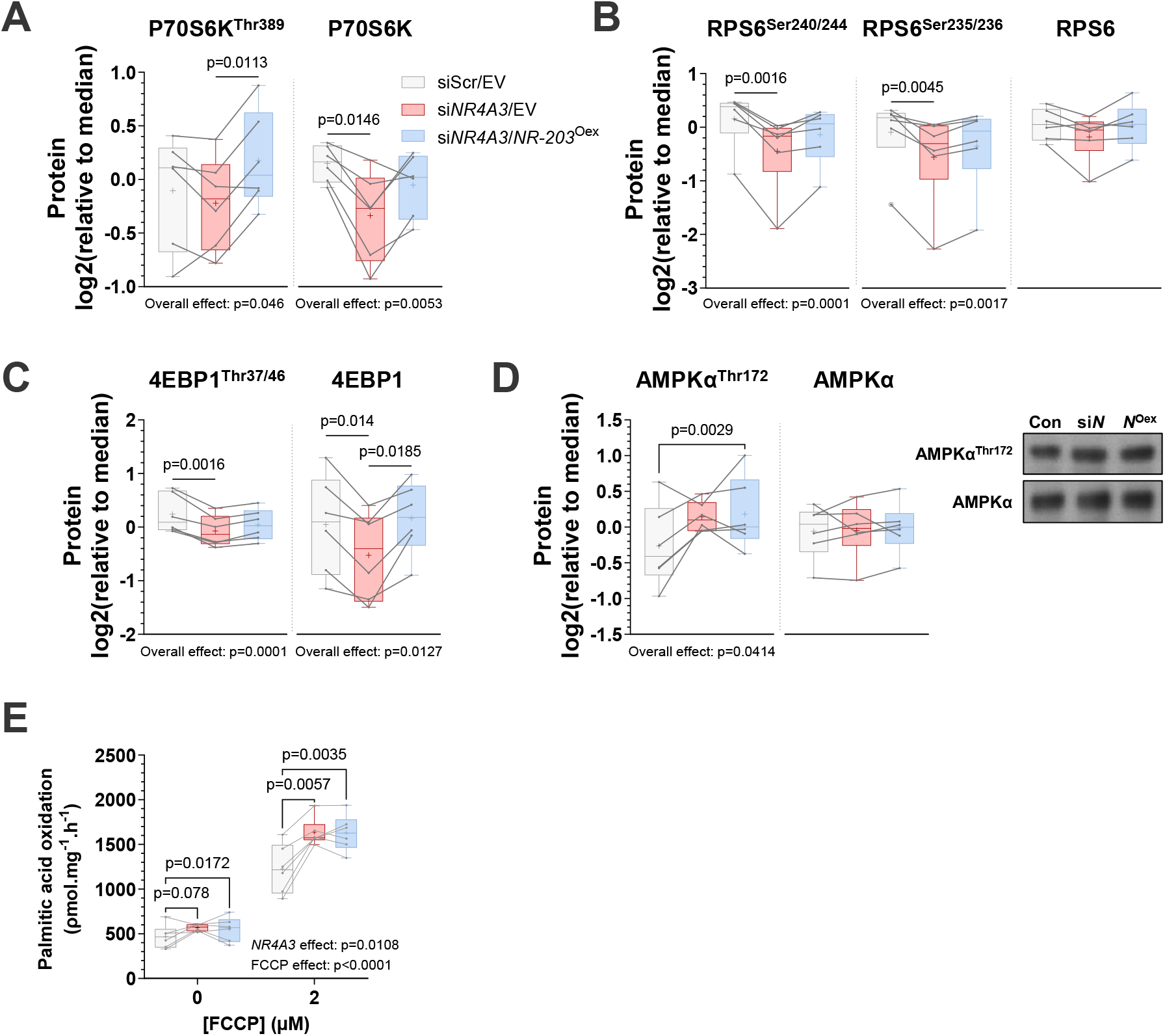
Reconstitution of the canonical *NR4A3* isoform after global NR4A3-depletion restores mTORC1 signalling but does not ameliorate AMPKα phosphorylation or fatty acid oxidation. Primary skeletal muscle cells were depleted of all *NR4A3* transcripts using siRNA and then transduced with an empty vector control plasmid or a plasmid containing variant *NR4A3-203* as described in main **Figure 6A. A-C**. Immunoblot quantification of total protein levels and phosphorylation events in the P70S6K-RPS6 and 4EBP1 branches of mTORC1 signalling. **D**. Representative immunoblot and quantification of total and phosphorylated AMPKα. All immunoblot analyses are n = 6, and either 1-way ANOVA with Tukey correction or Friedman test with Dunn’s correction. **E**. Rates of radiolabelled palmitic acid oxidation under basal and (2 µM) FCCP-stimulated conditions over 4 h. n = 6, 2-way ANOVA (overexpression x FCCP) with Fisher’s LSD post-test. Overall statistical model effects are stated in figures.

**Supplementary Figure 6.**
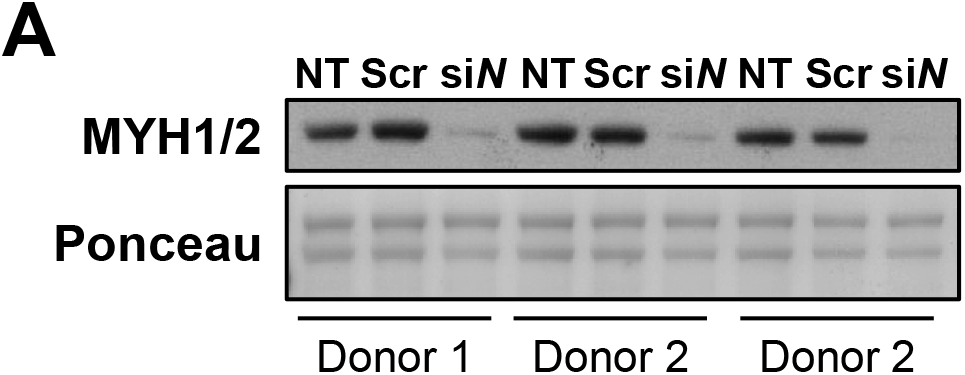
The effect of *NR4A3* silencing on contractile proteins is specific to siRNA targeting *NR4A3* transcripts. Primary skeletal muscle cells were exposed to a control scramble sequence (Scr) or a silencing RNA targeting *NR4A3* (si*N*). **A**. Representative immunoblot of myosin heavy chain isoforms IIX (MYH1) and IIA (MYH2) from three independent donors. ‘NT’ = non-transfected condition.

## Notes

### Competing Interest Statement

The authors have declared no competing interest.

### Summary of Updates

Updated experiments and figures based on feedback from other contributors to the study.

## References

Abdelmoez AM, Sardon Puig L, Smith JAB, Gabriel BM, Savikj M, Dollet L, Chibalin AV, Krook A, Zierath JR, Pillon NJ (2020) Comparative profiling of skeletal muscle models reveals heterogeneity of transcriptome and metabolism. Am J Physiol Cell Physiol 318: C615–C626

Amoasii L, Sanchez-Ortiz E, Fujikawa T, Elmquist JK, Bassel-Duby R, Olson EN (2019) NURR1 activation in skeletal muscle controls systemic energy homeostasis. Proc Natl Acad Sci U S A 116: 11299–11308

Baehr LM, Hughes DC, Lynch SA, Van Haver D, Maia TM, Marshall AG, Radoshevich L, Impens F, Waddell DS, Bodine SC (2021) Identification of the MuRF1 Skeletal Muscle Ubiquitylome Through Quantitative Proteomics. Function (Oxf) 2: zqab029

Bergouignan A, Antoun E, Momken I, Schoeller DA, Gauquelin-Koch G, Simon C, Blanc S (2013) Effect of contrasted levels of habitual physical activity on metabolic flexibility. J Appl Physiol (1985) 114: 371–379

Blum JE, Gheller BJ, Benvie A, Field MS, Panizza E, Vacanti NM, Berry D, Thalacker-Mercer A (2021) Pyruvate Kinase M2 Supports Muscle Progenitor Cell Proliferation but Is Dispensable for Skeletal Muscle Regeneration after Injury. J Nutr 151: 3313–3328

Bollinger LM, Witczak CA, Houmard JA, Brault JJ (2014) SMAD3 augments FoxO3-induced MuRF-1 promoter activity in a DNA-binding-dependent manner. Am J Physiol Cell Physiol 307: C278–287

Breen L, Stokes KA, Churchward-Venne TA, Moore DR, Baker SK, Smith K, Atherton PJ, Phillips SM (2013) Two weeks of reduced activity decreases leg lean mass and induces “anabolic resistance” of myofibrillar protein synthesis in healthy elderly. J Clin Endocrinol Metab 98: 2604–2612

Brook MS, Stokes T, Gorissen SHM, Bass JJ, McGlory C, Cegielski J, Wilkinson DJ, Phillips BE, Smith K, Phillips SM et al (2022) Declines in muscle protein synthesis account for short-term muscle disuse atrophy in humans in the absence of increased muscle protein breakdown. J Cachexia Sarcopenia Muscle 13: 2005–2016

Chao LC, Wroblewski K, Ilkayeva OR, Stevens RD, Bain J, Meyer GA, Schenk S, Martinez L, Vergnes L, Narkar VA et al (2012) Skeletal muscle Nur77 expression enhances oxidative metabolism and substrate utilization. J Lipid Res 53: 2610–2619

Eggelbusch M, Charlton BT, Bosutti A, Ganse B, Giakoumaki I, Grootemaat AE, Hendrickse PW, Jaspers Y, Kemp S, Kerkhoff TJ et al (2024) The impact of bed rest on human skeletal muscle metabolism. Cell Rep Med 5: 101372

Figueiredo VC, D’Souza RF, Van Pelt DW, Lawrence MM, Zeng N, Markworth JF, Poppitt SD, Miller BF, Mitchell CJ, McCarthy JJ et al (2021) Ribosome biogenesis and degradation regulate translational capacity during muscle disuse and reloading. J Cachexia Sarcopenia Muscle 12: 130–143

Gauthier-Coles G, Vennitti J, Zhang Z, Comb WC, Xing S, Javed K, Broer A, Broer S (2021) Quantitative modelling of amino acid transport and homeostasis in mammalian cells. Nat Commun 12: 5282

Goode JM, Pearen MA, Tuong ZK, Wang SC, Oh TG, Shao EX, Muscat GE (2016) The Nuclear Receptor, Nor-1, Induces the Physiological Responses Associated With Exercise. Mol Endocrinol 30: 660–676

Goodman CA, Mabrey DM, Frey JW, Miu MH, Schmidt EK, Pierre P, Hornberger TA (2011) Novel insights into the regulation of skeletal muscle protein synthesis as revealed by a new nonradioactive in vivo technique. FASEB J 25: 1028–1039

Gwinn DM, Shackelford DB, Egan DF, Mihaylova MM, Mery A, Vasquez DS, Turk BE, Shaw RJ (2008) AMPK phosphorylation of raptor mediates a metabolic checkpoint. Mol Cell 30: 214–226

Han JM, Jeong SJ, Park MC, Kim G, Kwon NH, Kim HK, Ha SH, Ryu SH, Kim S (2012) Leucyl-tRNA synthetase is an intracellular leucine sensor for the mTORC1-signaling pathway. Cell 149: 410–424

Hardy EJO, Inns TB, Hatt J, Doleman B, Bass JJ, Atherton PJ, Lund JN, Phillips BE (2022) The time course of disuse muscle atrophy of the lower limb in health and disease. J Cachexia Sarcopenia Muscle 13: 2616–2629

Inoki K, Zhu T, Guan KL (2003) TSC2 mediates cellular energy response to control cell growth and survival. Cell 115: 577–590

Kaiser MS, Milan G, Ham DJ, Lin S, Oliveri F, Chojnowska K, Tintignac LA, Mittal N, Zimmerli CE, Glass DJ et al (2022) Dual roles of mTORC1-dependent activation of the ubiquitin-proteasome system in muscle proteostasis. Commun Biol 5: 1141

Liang X, Liu L, Fu T, Zhou Q, Zhou D, Xiao L, Liu J, Kong Y, Xie H, Yi F et al (2016) Exercise Inducible Lactate Dehydrogenase B Regulates Mitochondrial Function in Skeletal Muscle. J Biol Chem 291: 25306–25318

Maira M, Martens C, Philips A, Drouin J (1999) Heterodimerization between members of the Nur subfamily of orphan nuclear receptors as a novel mechanism for gene activation. Mol Cell Biol 19: 7549–7557

Manini TM, Clark BC, Nalls MA, Goodpaster BH, Ploutz-Snyder LL, Harris TB (2007) Reduced physical activity increases intermuscular adipose tissue in healthy young adults. Am J Clin Nutr 85: 377–384

Massart J, Zierath JR, Chibalin AV (2014) A simple and rapid method to characterize lipid fate in skeletal muscle. BMC Res Notes 7: 391

Min K, Smuder AJ, Kwon OS, Kavazis AN, Szeto HH, Powers SK (2011) Mitochondrial-targeted antioxidants protect skeletal muscle against immobilization-induced muscle atrophy. J Appl Physiol (1985) 111: 1459–1466

Paez HG, Ferrandi PJ, Pitzer CR, Mohamed JS, Alway SE (2023) Loss of NOR-1 represses muscle metabolism through mTORC1-mediated signaling and mitochondrial gene expression in C2C12 myotubes. FASEB J 37: e23050

Pattamaprapanont P, Garde C, Fabre O, Barres R (2016) Muscle Contraction Induces Acute Hydroxymethylation of the Exercise-Responsive Gene Nr4a3. Front Endocrinol (Lausanne) 7: 165

Pavis GF, Abdelrahman DR, Murton AJ, Wall BT, Stephens FB, Dirks ML (2023) Short-term disuse does not affect postabsorptive or postprandial muscle protein fractional breakdown rates. J Cachexia Sarcopenia Muscle 14: 2064–2075

Pearen MA, Eriksson NA, Fitzsimmons RL, Goode JM, Martel N, Andrikopoulos S, Muscat GE (2012) The nuclear receptor, Nor-1, markedly increases type II oxidative muscle fibers and resistance to fatigue. Mol Endocrinol 26: 372–384

Pearen MA, Myers SA, Raichur S, Ryall JG, Lynch GS, Muscat GE (2008) The orphan nuclear receptor, NOR-1, a target of beta-adrenergic signaling, regulates gene expression that controls oxidative metabolism in skeletal muscle. Endocrinology 149: 2853–2865

Pillon NJ, Gabriel BM, Dollet L, Smith JAB, Sardon Puig L, Botella J, Bishop DJ, Krook A, Zierath JR (2020) Transcriptomic profiling of skeletal muscle adaptations to exercise and inactivity. Nat Commun 11: 470

Pin F, Novinger LJ, Huot JR, Harris RA, Couch ME, O’Connell TM, Bonetto A (2019) PDK4 drives metabolic alterations and muscle atrophy in cancer cachexia. FASEB J 33: 7778–7790

Pohl C, Dikic I (2019) Cellular quality control by the ubiquitin-proteasome system and autophagy. Science 366: 818–822

Popov A, Smirnov E, Kovacik L, Raska O, Hagen G, Stixova L, Raska I (2013) Duration of the first steps of the human rRNA processing. Nucleus 4: 134–141

Ritchie ME, Phipson B, Wu D, Hu Y, Law CW, Shi W, Smyth GK (2015) limma powers differential expression analyses for RNA-sequencing and microarray studies. Nucleic Acids Res 43: e47

Saxton RA, Knockenhauer KE, Wolfson RL, Chantranupong L, Pacold ME, Wang T, Schwartz TU, Sabatini DM (2016) Structural basis for leucine sensing by the Sestrin2-mTORC1 pathway. Science 351: 53–58

Shad BJ, Thompson JL, Holwerda AM, Stocks B, Elhassan YS, Philp A, LJC Vanl, Wallis GA (2019) One Week of Step Reduction Lowers Myofibrillar Protein Synthesis Rates in Young Men. Med Sci Sports Exerc 51: 2125–2134

Shur NF, Simpson EJ, Crossland H, Constantin D, Cordon SM, Constantin-Teodosiu D, Stephens FB, Brook MS, Atherton PJ, Smith K et al (2024) Bed-rest and exercise remobilization: Concurrent adaptations in muscle glucose and protein metabolism. J Cachexia Sarcopenia Muscle 15: 603–614

Siripoksup P, Cao G, Cluntun AA, Maschek JA, Pearce Q, Brothwell MJ, Jeong MY, Eshima H, Ferrara PJ, Opurum PC et al (2024) Sedentary behavior in mice induces metabolic inflexibility by suppressing skeletal muscle pyruvate metabolism. J Clin Invest 134

Smith JAB, Murach KA, Dyar KA, Zierath JR (2023) Exercise metabolism and adaptation in skeletal muscle. Nat Rev Mol Cell Biol 24: 607–632

Stephenson EJ, Stepto NK, Koch LG, Britton SL, Hawley JA (2012) Divergent skeletal muscle respiratory capacities in rats artificially selected for high and low running ability: a role for Nor1? J Appl Physiol (1985) 113: 1403–1412

Talbert EE, Smuder AJ, Min K, Kwon OS, Szeto HH, Powers SK (2013) Immobilization-induced activation of key proteolytic systems in skeletal muscles is prevented by a mitochondria-targeted antioxidant. J Appl Physiol (1985) 115: 529–538

Tontonoz P, Cortez-Toledo O, Wroblewski K, Hong C, Lim L, Carranza R, Conneely O, Metzger D, Chao LC (2015) The orphan nuclear receptor Nur77 is a determinant of myofiber size and muscle mass in mice. Mol Cell Biol 35: 1125–1138

von Walden F (2019) Ribosome biogenesis in skeletal muscle: coordination of transcription and translation. J Appl Physiol (1985) 127: 591–598

Weijs PJ, Looijaard WG, Dekker IM, Stapel SN, Girbes AR, Oudemans-van Straaten HM, Beishuizen A (2014) Low skeletal muscle area is a risk factor for mortality in mechanically ventilated critically ill patients. Crit Care 18: R12

West DW, Baehr LM, Marcotte GR, Chason CM, Tolento L, Gomes AV, Bodine SC, Baar K (2016) Acute resistance exercise activates rapamycin-sensitive and - insensitive mechanisms that control translational activity and capacity in skeletal muscle. J Physiol 594: 453–468

Wilson TE, Fahrner TJ, Johnston M, Milbrandt J (1991) Identification of the DNA binding site for NGFI-B by genetic selection in yeast. Science 252: 1296–1300

Wu T, Hu E, Xu S, Chen M, Guo P, Dai Z, Feng T, Zhou L, Tang W, Zhan L et al (2021) clusterProfiler 4.0: A universal enrichment tool for interpreting omics data. Innovation (Camb) 2: 100141

Yerrakalva D, Hajna S, Suhrcke M, Wijndaele K, Westgate K, Khaw KT, Wareham N, Brage S, Griffin S (2023) Associations between change in physical activity and sedentary time and health-related quality of life in older english adults: the EPIC-Norfolk cohort study. Health Qual Life Outcomes 21: 60

You JS, McNally RM, Jacobs BL, Privett RE, Gundermann DM, Lin KH, Steinert ND, Goodman CA, Hornberger TA (2019) The role of raptor in the mechanical load-induced regulation of mTOR signaling, protein synthesis, and skeletal muscle hypertrophy. FASEB J 33: 4021–4034

Yuan S, Li X, Liu Q, Wang Z, Jiang X, Burgess S, Larsson SC (2023) Physical Activity, Sedentary Behavior, and Type 2 Diabetes: Mendelian Randomization Analysis. J Endocr Soc 7: bvad090

